# Spatially Resolved In Silico Modeling of NKG2D Signaling Kinetics Reveals Key role of NKG2D and Vav1 Co-clustering in Generating Natural Killer cell Activation

**DOI:** 10.1101/2021.08.24.457567

**Authors:** Rajdeep Kaur Grewal, Jayajit Das

## Abstract

Natural Killer (NK) cells provide key resistance against viral infections and tumors. A diverse set of activating and inhibitory NK cell receptors (NKRs) interact with cognate ligands presented by target host cells, where integration of dueling signals initiated by the ligand-NKR interactions determines NK cell activation or tolerance. Imaging experiments over decades have shown micron and sub-micron scale spatial clustering of activating and inhibitory NKRs. The mechanistic roles of these clusters in affecting downstream signaling and activation are often unclear. To this end, we developed a predictive in silico framework by combining spatially resolved mechanistic agent based modeling, published TIRF imaging data, and parameter estimation to determine mechanisms by which formation and spatial movements of activating NKG2D microclusters affect early time NKG2D signaling kinetics in a human cell line NKL. We show co-clustering of NKG2D and the guanosine nucleotide exchange factor Vav1 in NKG2D microclusters plays a dominant role over ligand (ULBP3) rebinding in increasing production of phospho-Vav1(pVav1), an activation marker of early NKG2D signaling. The in silico model successfully predicts several scenarios of inhibition of NKG2D signaling and time course of NKG2D spatial clustering over a short (∼3 min) interval. Modeling shows the presence of a spatial positive feedback relating formation and centripetal movements of NKG2D microclusters, and pVav1 production offers flexibility towards suppression of activating signals by inhibitory KIR ligands organized in inhomogeneous spatial patterns (e.g., a ring). Our in silico framework marks a major improvement in developing spatiotemporal signaling models with quantitatively estimated model parameters using imaging data.

## Introduction

Natural Killer (NK) cells are lymphocytes of our innate immune system which provide important immune protection against viral infection and tumors ^1,2^. NK cells express a wide range of germ line encoded activating and inhibitory NK cell receptors (NKRs). In human NK cells, activating NKRs include NKG2D and killer Ig-like receptor (KIR) KIR2DS1, and inhibitory NKRs include inhibitory KIRs such as KIR2DL1, and KIR2DL2. NKRs bind to cognate ligands expressed by target cells initiating biochemical, physical, and mechanical modifications within NK cells that culminate into diverse NK cell responses ranging from a neutral response to lysis of target cells to secretion of cytokines^3^. Healthy host cells express polymorphic class I HLA molecules, cognate to a wide range of inhibitory NKRs, and generate tolerant NK cell responses. Whereas, tumor and infected cells downregulate expression of class I HLA molecules or upregulate expressions of ligands cognate to activating NKRs and tip the balance between activating and inhibitory signals toward NK cell activation. NKG2D is one of the best studied activating NKR and, in humans it binds to two families of ligands: one family (MICA and MICB) is akin to MHC class I and the other (ULBP1-3) is related to human proteins that bind to UL16 protein of human cytomegalovirus^4^. Tumor or infected host cells upregulate expressions of NKG2D ligands which contribute to lysis of these cells by NK cell cytotoxic responses^4^.

Spatial clustering of NKG2D has been well investigated in confocal^5^, total internal reflection fluorescence (TIRF)^6^, and super-resolution microscopy experiments^7^. Abeyweera et al. ^6^ using TIRF microscopy experiments reported formation of mobile and immobile microclusters of NKG2D in human NK cell line NKL stimulated by cognate ligand ULBP3 presented on a planar lipid bilayer supported by a glass coverslip. The NKG2D microclusters form at the interface between the NK cell and the lipid bilayer which is also known as the immunological synapse (IS). NKG2D microclusters, initially generated at the periphery of IS, migrate to the central region of the IS at later times, whereas NKG2D microclusters formed at the central region of the IS remain immobile. The mobility of NKG2D microclusters depends on actin remodeling as treatment by latrunculin, an actin depolymerizing agent, abrogated microcluster movements in the TIRF experiments. Additionally, confocal microscopy experiments show simultaneous localization of NKG2D receptors, Grb2, and Vav1 in the IS in NK cell line NKL stimulated by NKG2D ligand MICA^7^. Phosphorylation of Vav1 induces actin remodeling via activation of Rac GTPases, and thus can regulate motility of NKG2D microclusters.

Spatially resolved computational models have been successfully employed to glean mechanisms that underlie formation, motility, and function of spatial clusters of activating^8^ and inhibitory^9 10^ NKRs. Kaplan et al.^8^ developed a spatially resolved model to investigate hypotheses regarding signal integration of activating NKG2D and inhibitory human KIR2DL/mouse Ly49 receptors and concluded inhibitory NKRs locally suppress activating signals. Spatial in silico models describing cluster formation and signaling of inhibitory NKRs elucidated mechanisms giving rise peptide antagonism for inhibitory KIR2DL2/3^10^. However, the above in silico models do not quantitatively match or fit microcluster formation and movements of NKRs including NKG2D in silico with that observed in microscopy experiments. As we reason below, quantitative description of size and kinetics of NKG2D microclusters in in silico models could be important for gleaning mechanisms and generating improved model predictions regarding perturbation of signaling kinetics by drugs affecting NKG2D clustering (e.g., Cytochalasin D). (1) Formation of NKG2D microclusters can increase the production of phosphorylated Vav1 (pVav1) due to increase in the frequency of ligand (e.g., ULPB3) rebinding to NKG2D residing within microclusters, and/or increase in biochemical propensity of signaling reactions when NKG2D molecules are co-clustered with other signaling molecules (e.g., Vav1). (2) Increased production of pVav1 due to clustering of NKG2D can increase centripetal movements of NKG2D microclusters leading to higher spatial aggregation of NKG2D thereby further increasing pVav1 production. This chain of events constitutes a “spatial” positive feedback^11^. Therefore, quantitative magnitudes of sizes of NKG2D microclusters and spatial separations between these microclusters could be relevant for affecting downstream signaling.

Motivated by above reasoning and a potential interplay between signaling kinetics and spatial clustering, we developed a framework combining spatially resolved mechanistic in silico agent based model and published TIRF imaging data assaying spatiotemporal signaling kinetics in NKL cell lines stimulated via NKG2D and/or KIR2DL2 receptors. The agent based model developed here represents a major improvement over previous modeling efforts in the following aspects: (1) The model is able to quantitatively describe micron scale details of spatial clustering of activating and inhibitory NKRs. (2) A detailed parameter estimation of model parameters is carried out using spatial data. (3) The model includes interplay between spatial clustering and activating NKR signaling kinetics. We investigated three hypotheses (Model 1, Model 2, and Model 3) to determine the role of co-clustering of signaling protein Vav1 with NKG2D and show that co-clustering of Vav1 with NKG2D is required to produce increased pVav1 production due to the formation of NKG2D microclusters, and the presence of the spatial positive feedback allows for an efficient suppression of early time NKG2D signaling by heterogeneously distributed inhibitory HLA-C ligands on target cells.

### Model Development

We developed a spatially resolved agent based model involving activating NKG2D receptors, inhibitory KIR2DL2 receptors, cognate NKR ligands (ULBP3 and HLA-C), and signaling proteins: Src family kinases (SFKs), Vav1, and phosphatase SHP1. The NKRs and signaling proteins interact via different rules to describe membrane proximal signaling events in human NK cell line NKL. The model includes rules describing biochemical signaling reactions, spatial movements of signaling complexes, and regulation of the spatial movements by biochemical reactions. The rules in the model are designed to quantitatively capture several distinguishing features of spatiotemporal NK cell signaling kinetics observed in previous imaging experiments, namely, a) formation of mobile and immobile NKG2D microclusters upon NKG2D stimulation^6^, b) centripetal movements of the mobile NKG2D microclusters from the periphery to the central region of the IS^6,12^, c) decrease in the velocity of mobile NKG2D microclusters as those move closer to the central region of the IS^6^, and, d) random movements of NKG2D microclusters in the central region of the IS^6^.

Our agent based model describes spatiotemporal membrane proximal NK cell signaling in a quasi three-dimensional simulation box representing the interface between NK cell membrane and the plasma membrane of target cell or the lipid bilayer in a TIRF experiment at the IS (Figure 1). The simulation box of area (A=15µm × 15µm) and thickness *z* is discretized into chambers of volume *l*_0_× *l*_0_×*z* where, *z*=*l*_0_ or *z*=2*l*_0_ (*l*_0_ = 0.5µm) for molecules residing in the plasma membrane or the cytosol, respectively. The following spatiotemporal processes occur in the models. (i) *<u>Biochemical signaling reactions</u>*: NKG2D receptors bind to cognate ligands (ULBP3) to form NKG2D-ULPB3 complexes. NKG2D homodimers are associated with a pair of DAP10 homodimers in NK cells. The tyrosine residues in two YINM motifs in DAP10 homodimer are phosphorylated by Src family kinases upon ULBP3 binding^13^. Thus, the DAP10 molecules associated with NKG2D homodimers can be in a variety of partially phosphorylated states where one, two or three of the total four tyrosine residues phosphorylated. We approximate the partially and fully phosphorylated states of the two DAP10 homodimers by two states, unphosphorylated or fully phosphorylated, to reduce the number of agents in the model. In the model, a kinase molecule (SFK) represents SFKs that phosphorylate DAP10 upon formation of NKG2D-ULBP3 complex. In NK cells, phosphorylated DAP10 becomes available to bind Grb2-Vav1 complex^1,14^. In the model, these reactions are approximated by binding of Vav1 to fully phosphorylated DAP10 where Grb2 is not included explicitly. DAP10 bound Vav1 is phosphorylated by the SFK in the model. Vav1 phosphorylation is an important event during early time NK cell signaling as pVav1 leads to actin polymerization and degranulation in NK cells^2,14^. Inhibitory KIR2DL2 receptors bind to cognate ligands (HLA-C) and tyrosine residues in immunoreceptor tyrosine based inhibitory motifs (ITIMs) associated with cytoplasmic part of KIR2DL2 are phosphorylated by the SFKs^15^. The model assumes two states of ITIM phosphorylation (unphosphorylated and fully phosphorylated) and the phosphatase SHP-1 binds to fully phosphorylated ITIMs. ITIM bound SHP-1 dephosphorylates pVav1 via enzymatic reactions^15^. Unbinding of ligands from cognate receptors (NKG2D or KIR2DL2) dissociates the signaling complexes completely in the model – this step is included to implement kinetic proofreading^16,17^. In addition, there are first order dephosphorylation reactions for pVav1 representing dephosphorylation of phospho-tyrosines by phosphatases other than SHP1. (ii) *<u>Spatial movements</u>*: We modeled movements of NKG2D-ULBP3 complexes for formation of NKG2D microclusters and centripetal movements of NKG2D microclusters towards the center of the IS. These movements are assumed in the model to be dependent on actin remodeling which is regulated by signaling products such as pVav1^18^. The above movements are implemented by nearest neighbor hops occurring with specific probabilities. The velocity of the microclusters decreases as those move closer to the center. In addition, unbound molecules of receptor, ligand, and signaling proteins perform diffusive random movements in the model.

**Figure 1.**
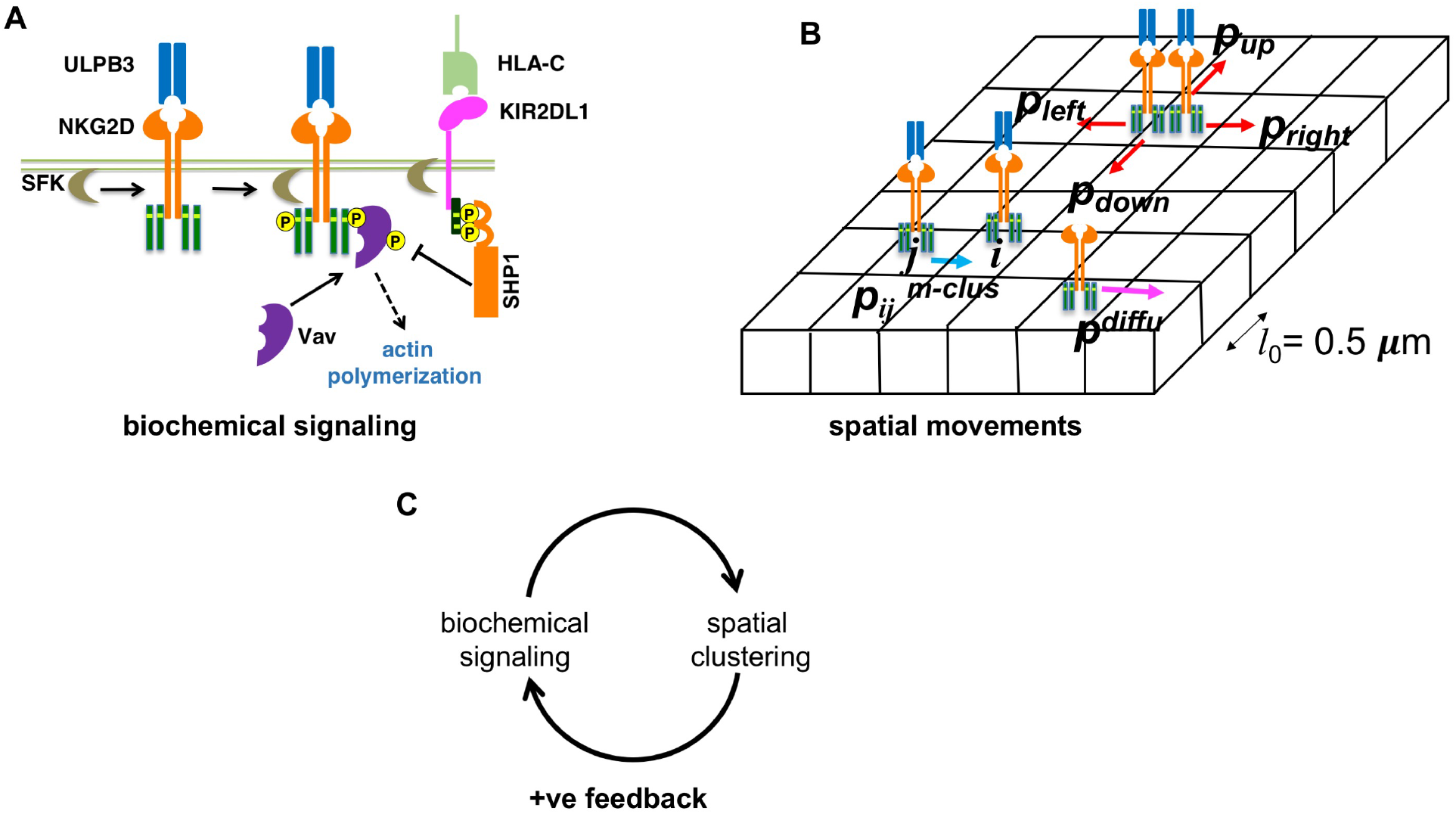
Schematic depiction of the agent based models. **(A)** Shows biochemical signaling reactions considered in the models. The reactions and their propensities are shown in Table 1. **(B)** Shows spatial movements considered in the agent based models. The simulation box is divided into chambers of volume *l*_0_ × *l*_0_ × *z*. A ULBP3 bound NKG2D complex at chamber *i* moves to its nearest neighbor chamber *j* with a probability *p*^(m-clus)^_*ij*_. *p*^(m-clus)^_*ij*_ depends on the pVav1 numbers in chamber *i* and the four nearest neighbor chambers. ULPB3 bound NKG2D complexes in a chamber hop to the nearest neighbor chambers with probabilities *p*_left_, *p*_right_, *p*_down_ and *p*_up_ to implement centripetal movements (see main text). Free protein (receptors, kinases, phosphatases, Vav1/pVav1) molecules hop to next nearest neighbor chambers with probability *p*^diffu^ implementing diffusive moves. (C) The biochemical signaling induces spatial clustering of NKG2D which in turn increases biochemical signaling-this represents a spatial positive feedback in the models.

Further details are provided in Table 1, Materials and Methods section, and the Supplementary Material. (iii) *<u>Interplay between spatial movements and signaling reactions</u>*: The probabilities of hops generating NKG2D microclusters and centripetal movements of NKG2D microclusters depend on the local and total number of pVav1 in the simulation box, respectively. The rules capture the interplay between biochemical signaling reactions and regulation of specific spatial movements by signaling reactions which in turn affects signaling reactions. For NKG2D signaling the above interplay represents a positive feedback^11^.

**Table 1:**
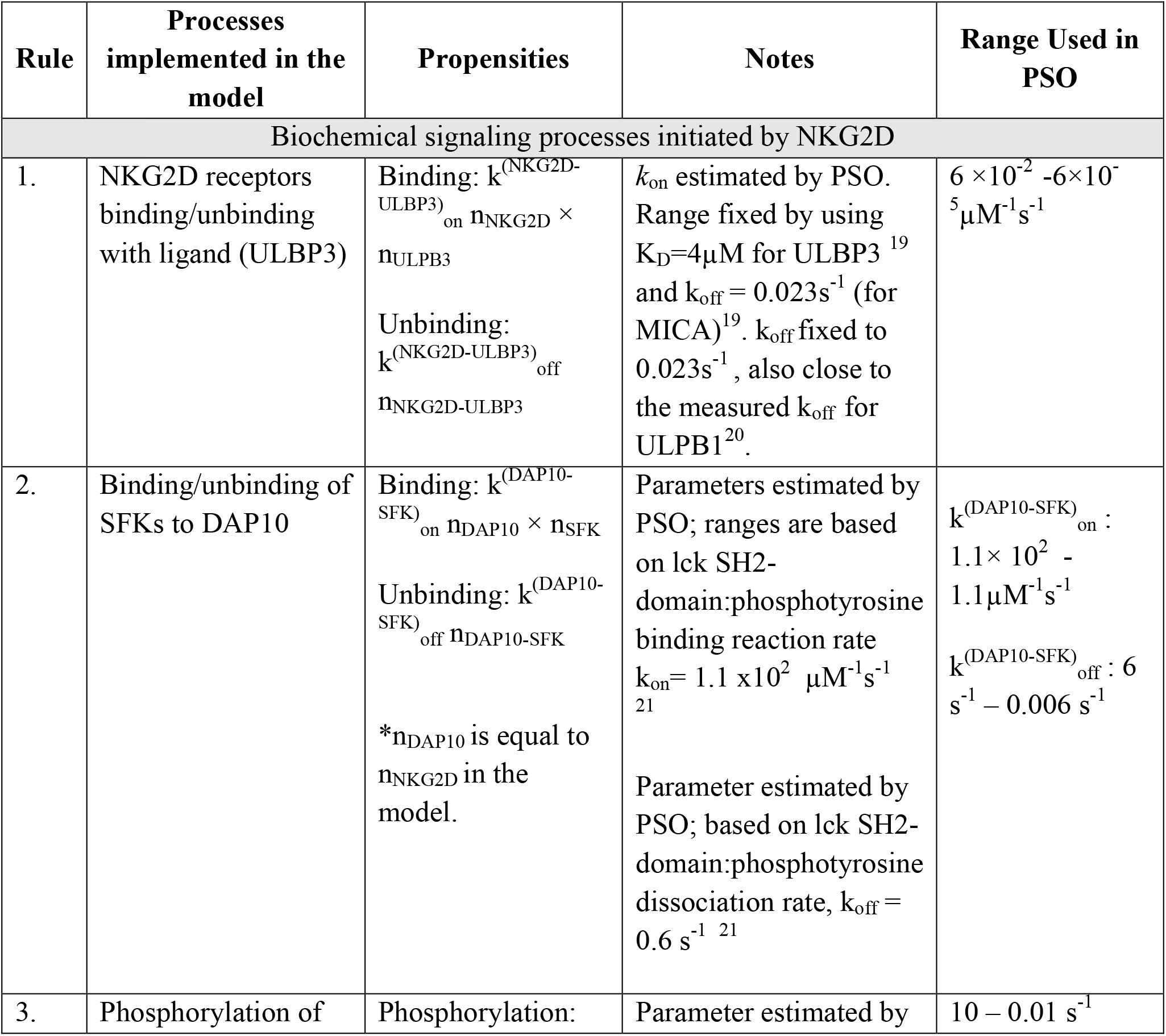
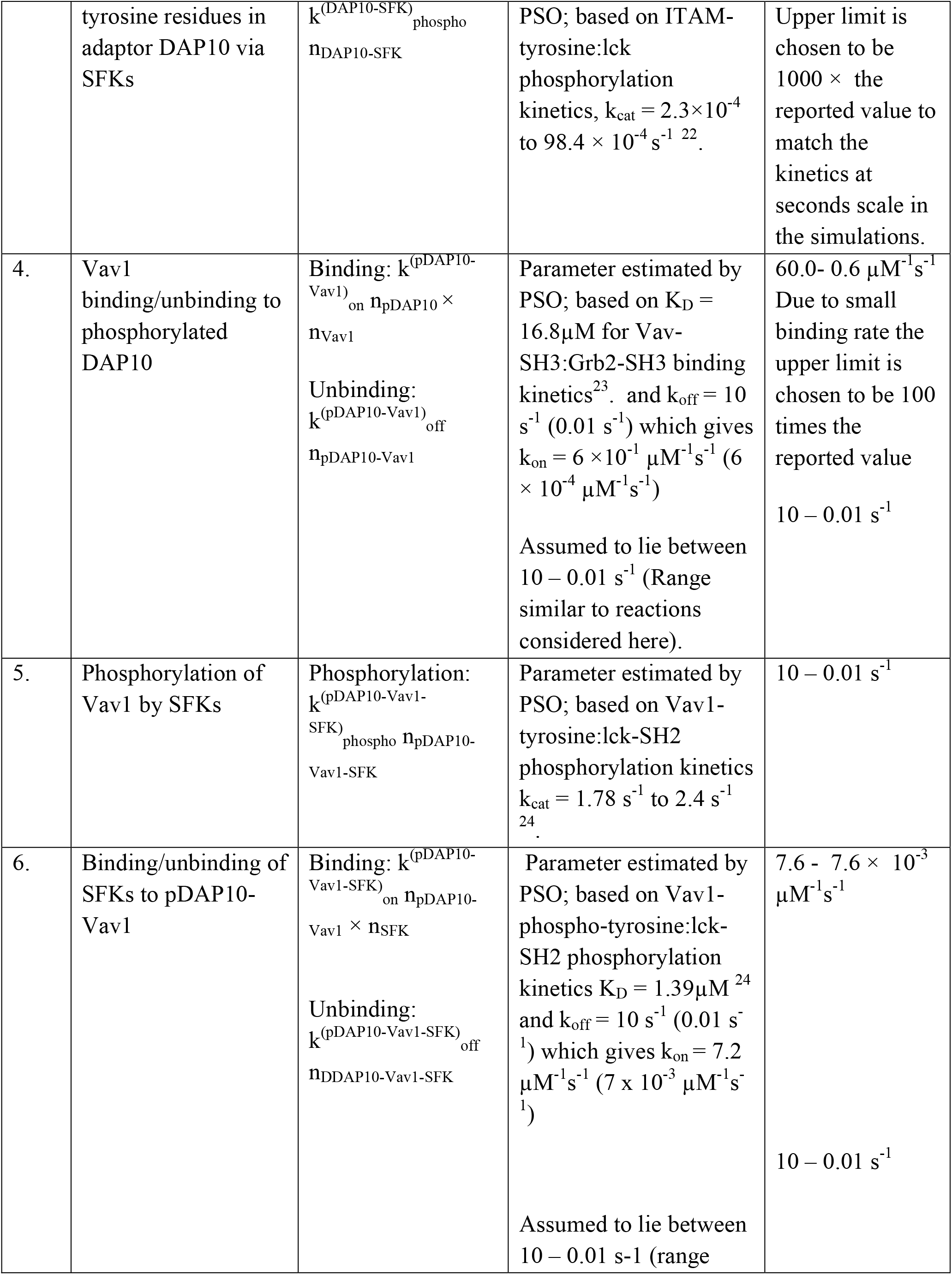
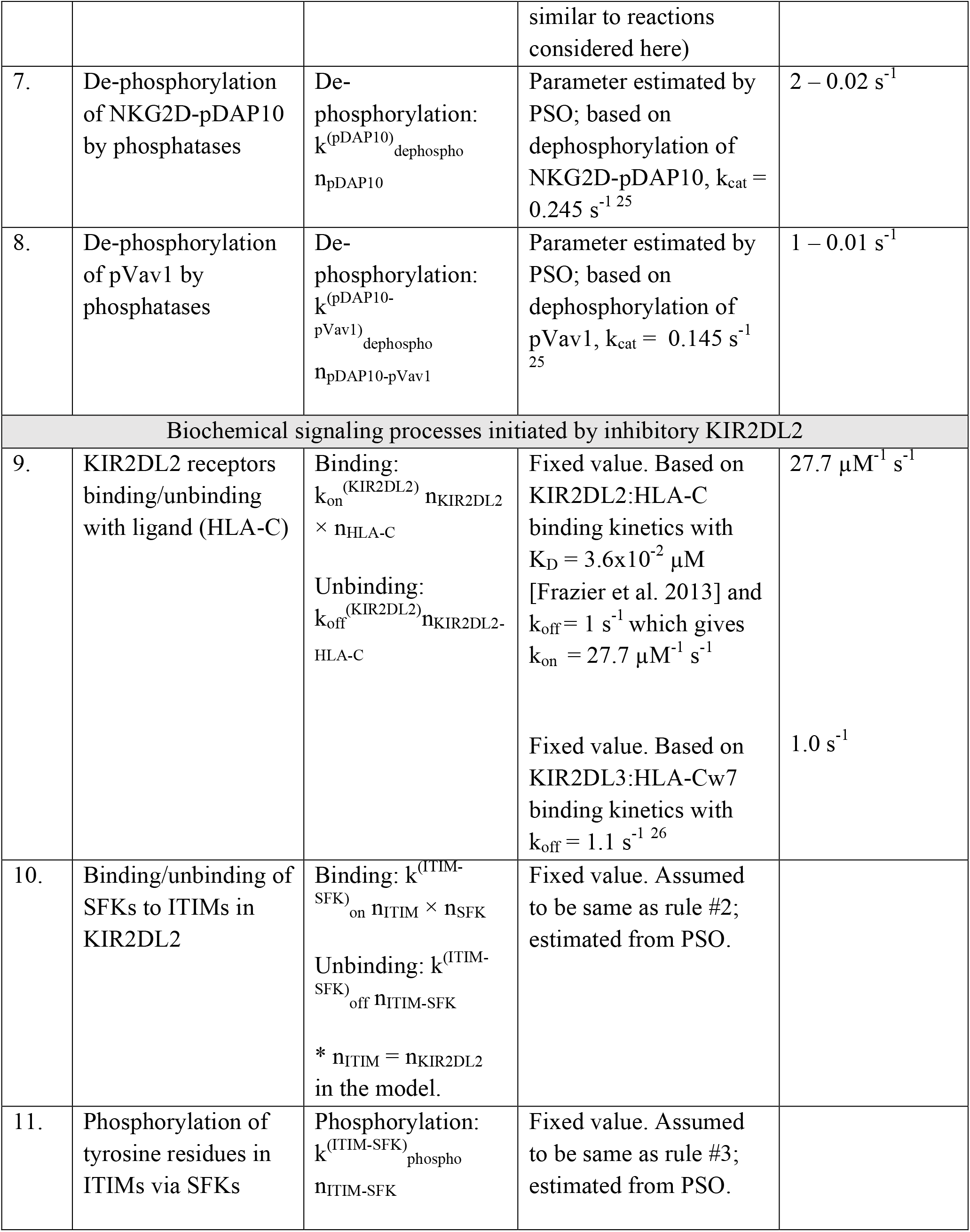
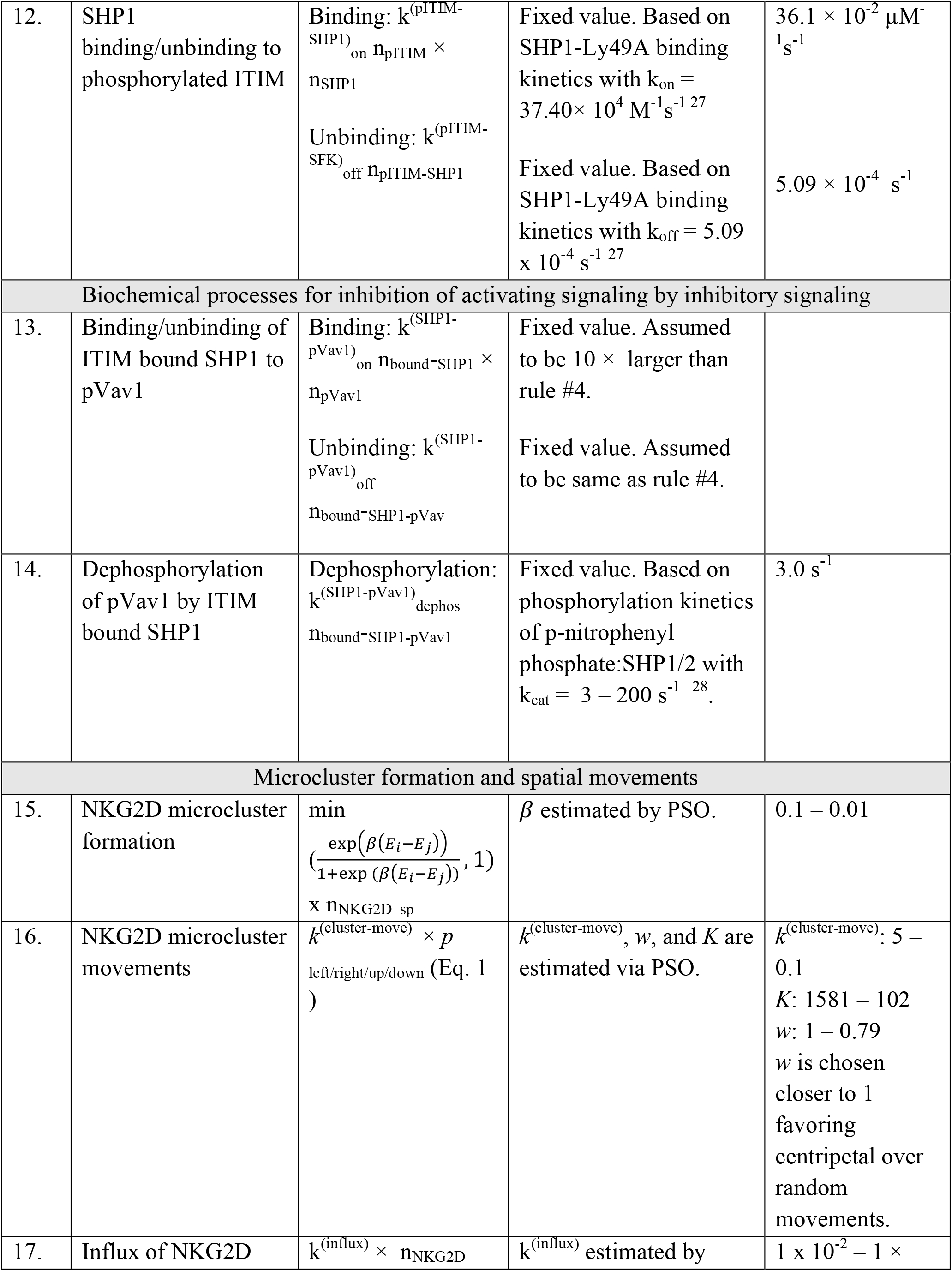
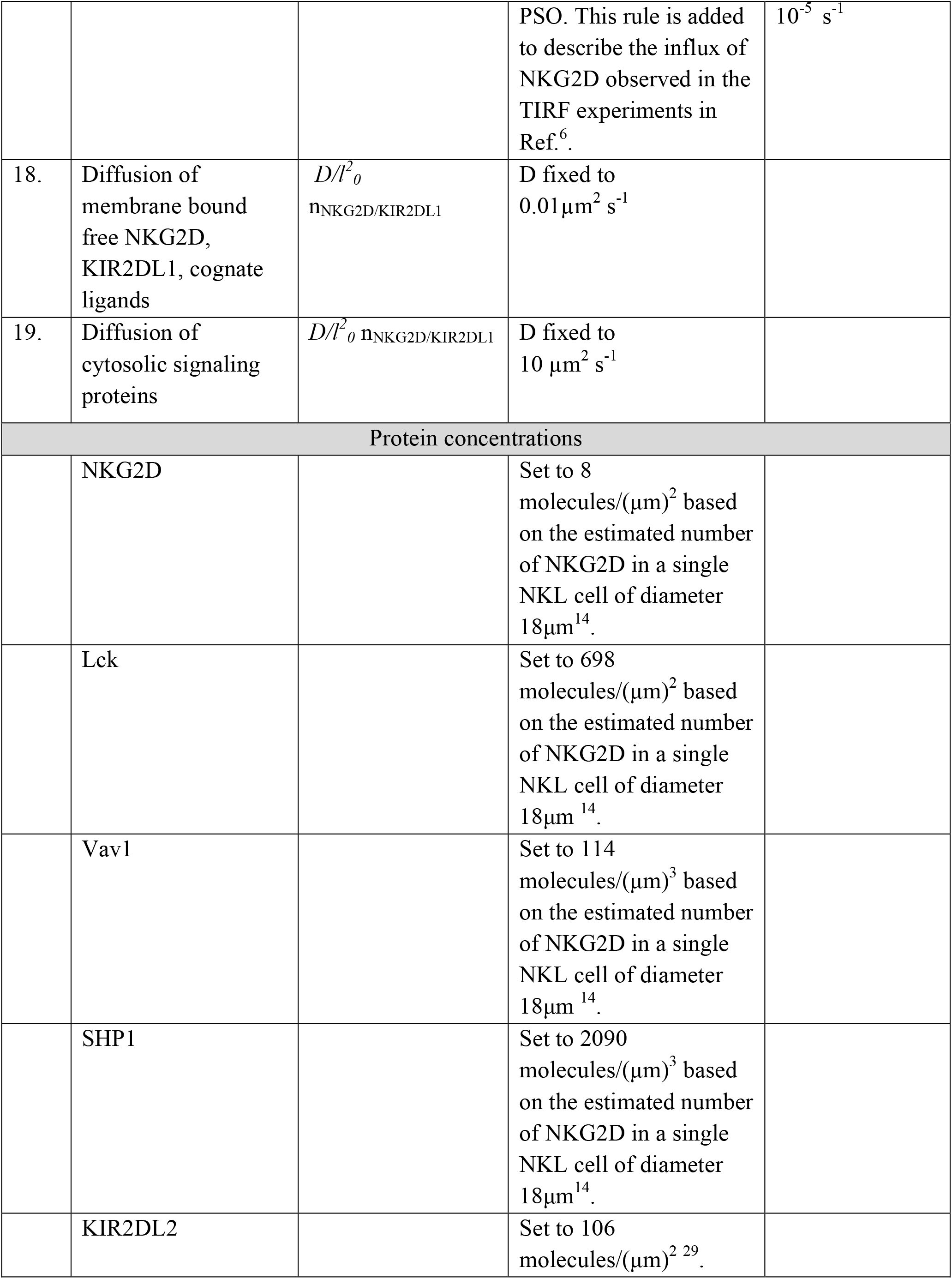
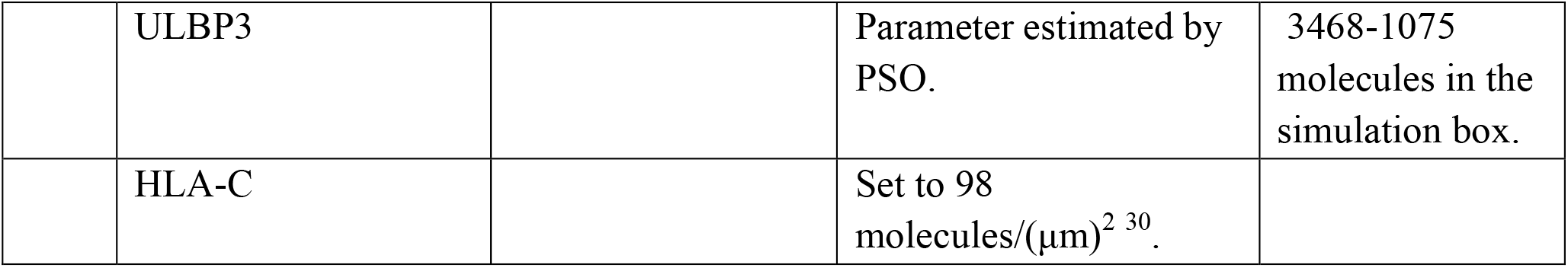
List of processes and parameter values used in the agent based model.

#### Hypotheses considered

We considered three hypotheses, encoded in models, Model 1, Model 2, and Model 3, which probe functional consequences of different types of NKG2D clustering and mechanisms of interplay between NKG2D microclusters and biochemical signaling reactions. In Model 1, mobile NKG2D microclusters and signaling molecules bound to NKG2D move toward the central region of the IS. In Model 2, NKG2D and Vav1 molecules residing in the proximity of the plasma membrane move simultaneously toward the central region of the IS. This Vav1 species could potentially represent Vav1 molecules that are recruited to the plasma membrane via SOS1^31^. Consequently, there is a strong co-clustering of NKG2D and Vav1 in Model 2 but a weak co-clustering of NKG2D and other signaling molecules in Model 1. In Model 3, we studied outcomes of the absence of the positive feedback between NKG2D microcluster movement and signaling reactions. In Model 3, the rate of centripetal movements of mobile NKG2D microclusters is independent of pVav1 abundances. Model 3 contains co-clustering of NKG2D and Vav1 as in Model 2. The models are summarized in Table 2.

**Table 2:** List of the agent based models.

|  | Biochemical signaling reactions | Formation of NKG2D microclusters | Centripetal movement of NKG2D microclusters | Co-clustering of NKG2D and Vav1 | pVav1 dependent centripetal movements of NKG2D micro clusters |
| --- | --- | --- | --- | --- | --- |
| **Model 1** | Yes | Yes | Yes | No | Yes |
| **Model 2** | Yes | Yes | Yes | Yes | Yes |
| **Model 3** | Yes | Yes | Yes | Yes | No |

### Model Simulation and Parameter estimation

The simulations are carried out in a quasi-three-dimensional simulation box representing a thin junction between NK cell and the supported lipid bilayer in TIRF experiments. The simulation box has an area of 15 × 15 µm^2^ and a depth of *z*, and is divided into small cubic chambers of size (*l*_0_ × *l*_0_ × *z*), where *l*_0_=0.5µm and *z*= *l*_0_ (for molecules residing in membrane) or 2*l*_0_ (for cytosolic molecules). The molecules are well-mixed in each chamber and molecules in a chamber hop to next nearest neighboring chambers with specific rates to produce diffusive or centripetal movements, or movements leading to microcluster formation. The kinetics of the system is simulated using a kinetic Monte Carlo (kMC) approach, which is implemented via a freely available simulator known as SPPARKS (https://spparks.sandia.gov/). The kMC simulation includes intrinsic noise fluctuations in biochemical reactions as well as in spatial movements.

The list of the processes, and their propensities are listed in Table 1. The copy numbers of most of the signaling proteins in the simulation box are estimated using available measured concentrations for NKLs (Table 1). However, values of many of the model parameters in the cellular environment are unknown, and we estimated these parameters by a parameter fitting scheme that reproduced the spatial pattern of NKG2D receptors measured in TIRF experiments as reported in Ref. ^6^. The spatial patterns of NKG2D in our simulation and TIRF imaging data are quantified using mean values, variances, and a two-point correlation function computed from density profiles for NKG2D. Similar variables are widely used in statistical physics^32^ to quantify spatial patterns. Euclidean distance between dimensionless forms of the above variables in TIRF images and the agent based model is used to create a cost function which is minimized by particle swarm optimization (PSO) to estimate model parameters. Details regarding our parameter estimation scheme are provided in Materials and Methods section and the Supplementary Material.

## Results

### 1. Multiple models quantitatively describe kinetics of NKG2D microclusters

Spatiotemporal signaling kinetics of NKG2D and associated signaling proteins was simulated in Model 1 and Model 2. NKG2D, ULBP3, SFK, and Vav1 molecules were distributed homogeneously in space in the simulation box at the beginning of the simulation at t=0. Binding of NKG2D and ULBP3 initiate a series of biochemical signaling reactions (Table 1) leading to phosphorylation of Vav1 in the simulations. The production of pVav1 molecules induce formation of NKG2D microclusters and centripetal movements of the microclusters in the simulations (Movies S1 and S2). Both models fit spatial distribution of NKG2D in TIRF experiments at t=1min, in particular at length scales ≥ 1µm, reasonably well (Figure 2). The variances and the two-point correlation functions calculated from spatial distributions of NKG2D in model simulations agreed well with that of TIRF imaging data (Figure 2 and Figure S1). The estimated best-fit parameters for the models are shown in Table 3. Many parameter values show order of magnitude differences between Model 1 and Model 2, e.g., binding-unbinding rates of SFK to DAP10 or of Vav1 to pDAP10. The reason for this difference can be explained in the following manner. In Model 2, co-clustering of Vav1 and NKG2D increases reaction propensities that affect formation and motility of NKG2D microclusters. Therefore, in order to compensate for the absence of the increase in reaction propensities due to Vav1-NKG2D co-clustering in Model 1, different values for reaction rates (e.g., binding/unbinding rate of Vav1 to pDAP10) regulating microcluster kinetics are chosen as optimal parameter values.

**Table 3:**
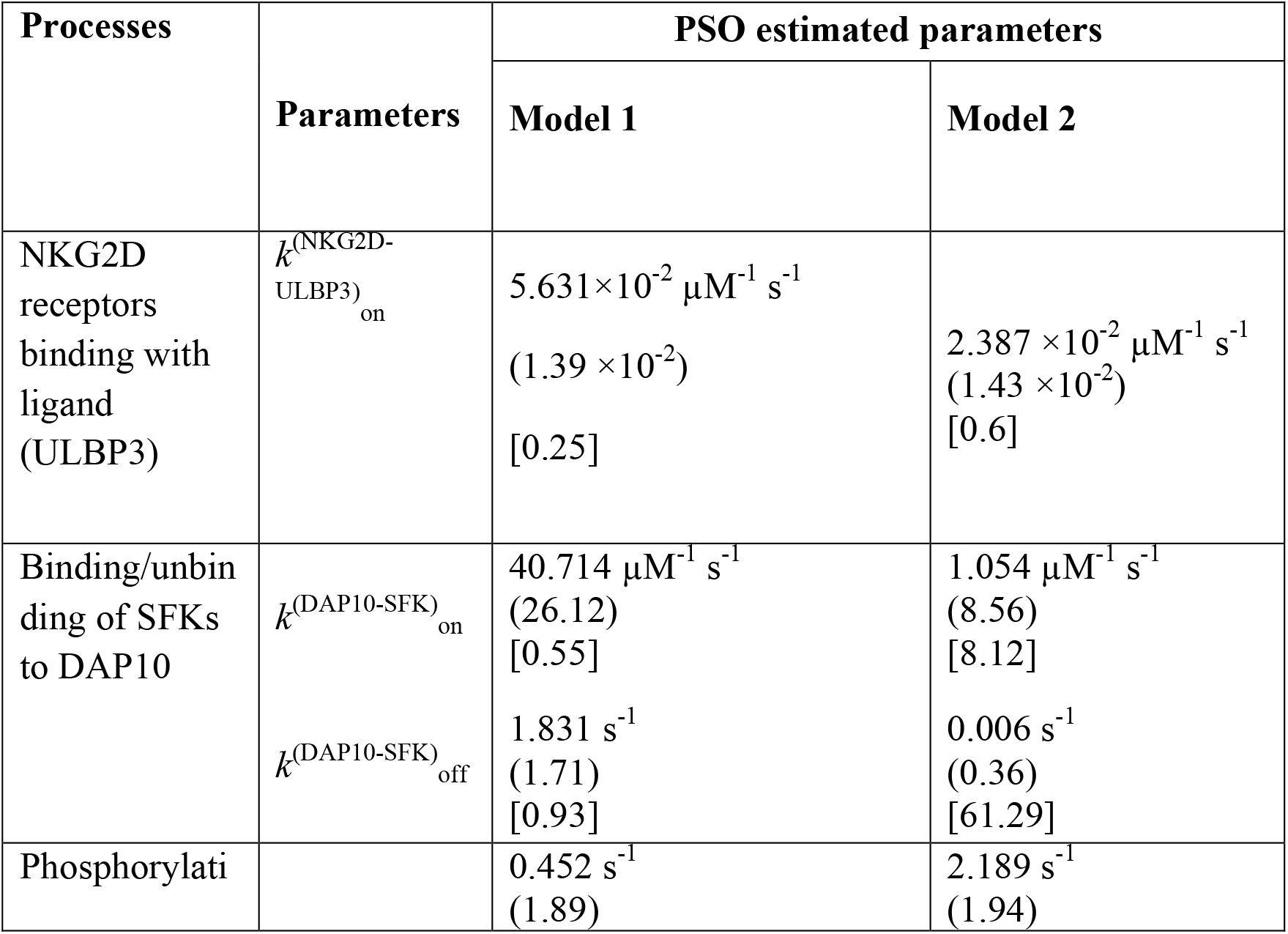
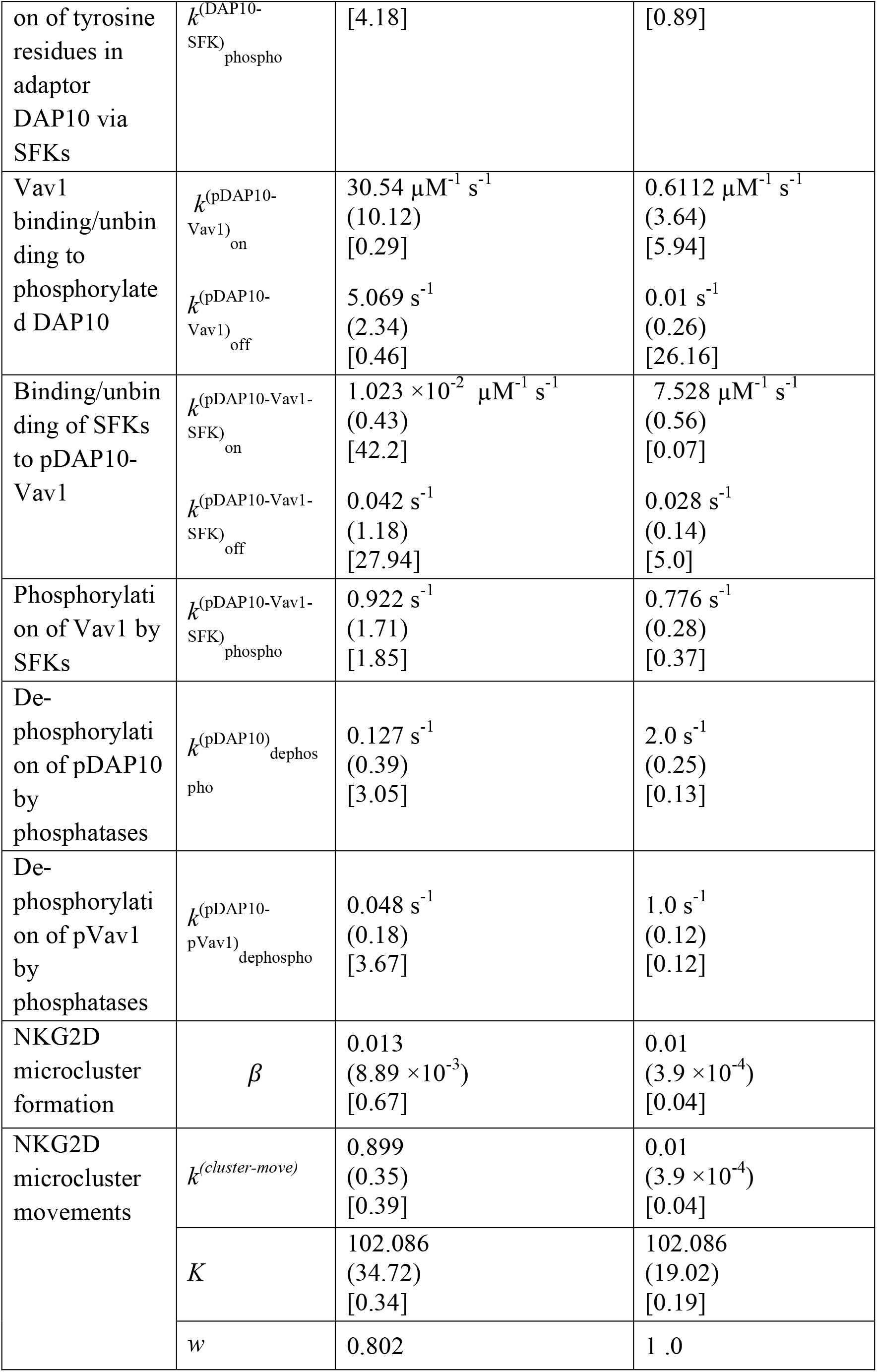
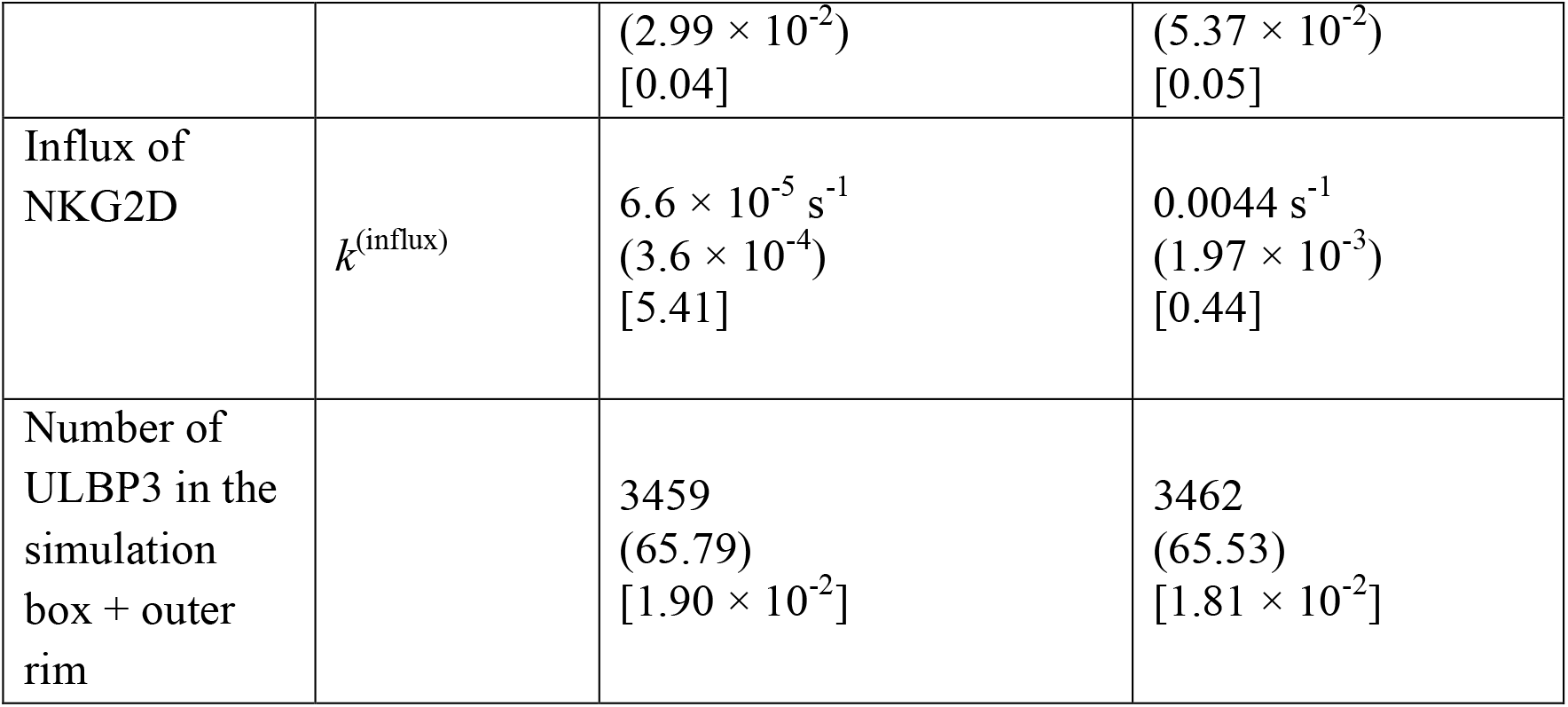
List of parameter values estimated from PSO. Values within () and [] correspond to the standard deviation (s.d.) (details in Materials and Methods section) and to the ratio PSO s.d./(PSO estimated value), respectively.

| Processes | Parameters | PSO estimated parameters |  |
| --- | --- | --- | --- |
|  |  | Model 1 | Model 2 |
| NKG2D receptors binding with ligand (ULBP3) | $k_{\text{ULBP3}}^{(\text{NKG2D})}_{\text{on}}$ | $5.631 \times 10^{-2} \mu\text{M}^{-1} \text{s}^{-1}$<br>($1.39 \times 10^{-2}$)<br>[0.25] | $2.387 \times 10^{-2} \mu\text{M}^{-1} \text{s}^{-1}$<br>($1.43 \times 10^{-2}$)<br>[0.6] |
| Binding/unbinding of SFKs to DAP10 | $k_{\text{on}}^{(\text{DAP10-SFK})}$ | $40.714 \mu\text{M}^{-1} \text{s}^{-1}$<br>(26.12)<br>[0.55] | $1.054 \mu\text{M}^{-1} \text{s}^{-1}$<br>(8.56)<br>[8.12] |
| | $k_{\text{off}}^{(\text{DAP10-SFK})}$ | $1.831 \text{s}^{-1}$<br>(1.71)<br>[0.93] | $0.006 \text{s}^{-1}$<br>(0.36)<br>[61.29] |
| Phosphorylation | | $0.452 \text{s}^{-1}$<br>(1.89) | $2.189 \text{s}^{-1}$<br>(1.94) |
| on of tyrosine residues in adaptor DAP10 via SFKs | $k^{(\text{DAP10-SFK})}_{\text{phospho}}$ | [4.18] | [0.89] |
| Vav1 binding/unbinding to phosphorylated DAP10 | $k^{(\text{pDAP10-Vav1})}_{\text{on}}$<br>$k^{(\text{pDAP10-Vav1})}_{\text{off}}$ | $30.54 \mu\text{M}^{-1} \text{s}^{-1}$<br>(10.12)<br>[0.29]<br>$5.069 \text{s}^{-1}$<br>(2.34)<br>[0.46] | $0.6112 \mu\text{M}^{-1} \text{s}^{-1}$<br>(3.64)<br>[5.94]<br>$0.01 \text{s}^{-1}$<br>(0.26)<br>[26.16] |
| Binding/unbinding of SFKs to pDAP10-Vav1 | $k^{(\text{pDAP10-Vav1-SFK})}_{\text{on}}$<br>$k^{(\text{pDAP10-Vav1-SFK})}_{\text{off}}$ | $1.023 \times 10^{-2} \mu\text{M}^{-1} \text{s}^{-1}$<br>(0.43)<br>[42.2]<br>$0.042 \text{s}^{-1}$<br>(1.18)<br>[27.94] | $7.528 \mu\text{M}^{-1} \text{s}^{-1}$<br>(0.56)<br>[0.07]<br>$0.028 \text{s}^{-1}$<br>(0.14)<br>[5.0] |
| Phosphorylation of Vav1 by SFKs | $k^{(\text{pDAP10-Vav1-SFK})}_{\text{phospho}}$ | $0.922 \text{s}^{-1}$<br>(1.71)<br>[1.85] | $0.776 \text{s}^{-1}$<br>(0.28)<br>[0.37] |
| De-phosphorylation of pDAP10 by phosphatases | $k^{(\text{pDAP10})}_{\text{dephospho}}$ | $0.127 \text{s}^{-1}$<br>(0.39)<br>[3.05] | $2.0 \text{s}^{-1}$<br>(0.25)<br>[0.13] |
| De-phosphorylation of pVav1 by phosphatases | $k^{(\text{pDAP10-pVav1})}_{\text{dephospho}}$ | $0.048 \text{s}^{-1}$<br>(0.18)<br>[3.67] | $1.0 \text{s}^{-1}$<br>(0.12)<br>[0.12] |
| NKG2D microcluster formation | $\beta$ | 0.013<br>($8.89 \times 10^{-3}$)<br>[0.67] | 0.01<br>($3.9 \times 10^{-4}$)<br>[0.04] |
| NKG2D microcluster movements | $k^{(\text{cluster-move})}$ | 0.899<br>(0.35)<br>[0.39] | 0.01<br>($3.9 \times 10^{-4}$)<br>[0.04] |
| | $K$ | 102.086<br>(34.72)<br>[0.34] | 102.086<br>(19.02)<br>[0.19] |
| | $w$ | 0.802 | 1.0 |
| | | $(2.99 \times 10^{-2})$<br>[0.04] | $(5.37 \times 10^{-2})$<br>[0.05] |
| Influx of NKG2D | $k^{(\text{influx})}$ | $6.6 \times 10^{-5} \text{ s}^{-1}$<br>$(3.6 \times 10^{-4})$<br>[5.41] | $0.0044 \text{ s}^{-1}$<br>$(1.97 \times 10^{-3})$<br>[0.44] |
| Number of ULBP3 in the simulation box + outer rim | | 3459<br>(65.79)<br>[$1.90 \times 10^{-2}$] | 3462<br>(65.53)<br>[$1.81 \times 10^{-2}$] |

**Figure 2.**
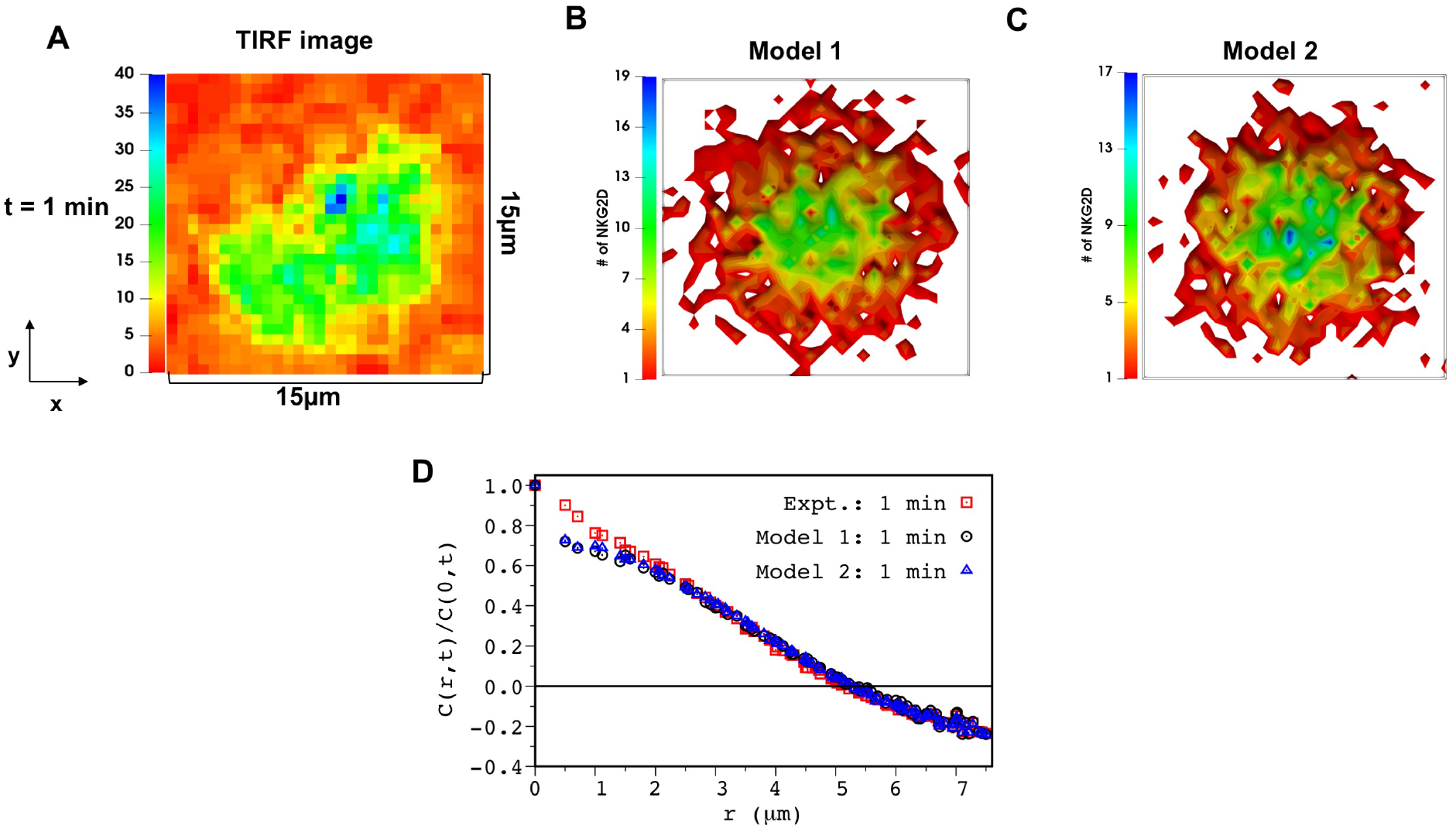
Model 1 and Model 2 captures the spatial clustering of NKG2D (DAP10) in TIRF experiments. **(A)** Shows a 2D fluorescence intensity of NKG2D-DAP10-mCherry in TIRF experiments in Ref. ^6^ at t=1min post stimulation by ULPB3. The above image shows a region of interest extracted from Figure S4 in Ref. ^6^ which is coarse-grained to match the minimum length scale (∼ 0.5 µm) of spatial resolution in our model (see Materials and Methods for details). **(B and C)** Shows spatial distribution of NKG2D in the x-y plane in the simulation box for our agent based model (B) Model 1 and (C) Model 2 at t=1 min post NKG2D stimulation. The parameters for the simulation are set at the best fit value from our PSO. The area of the region of interest in the image and simulation box is set to 15µm × 15µm. **(D)** Shows comparison between the two-point correlation function (C(r,t)/C(0,t) vs r) at t=1 min calculated from the image in (A) (red, empty square) and configurations for Model 1 (black, empty circle) and Model 2 (blue, empty triangle) shown in (B) and (C).

The computation of uncertainties in the estimated parameter values showed that about 2/3 of the total number of parameters are estimated well, e.g., standard deviation/estimated value < 3. The procedure for estimation of uncertainty is described in detail in the Materials and Methods section, and potential causes behind poor parameter estimations of 1/3 of the parameters are discussed in the Discussion section. Next, we assessed the utility of the parameter estimation scheme for generating predictions at future time points (Figure 3 and Figure S2) and providing mechanistic insights (next section). The best-fit models were used for predicting spatial patterns of NKG2D at later times t > 1 min. Both models predict spatial distribution of NKG2D in TIRF imaging data until t=3 min reasonably well (Figure 3 and Figure S2). However, model predictions deviate from the C(r,t) data calculated for the TIRF images at t=4 min (Figure S3). This disagreement is likely due to change in the organization of the NKG2D microclusters due to the spreading of the NK cells on the supported lipid bilayer which is not included in the agent based models.

**Figure 3.**
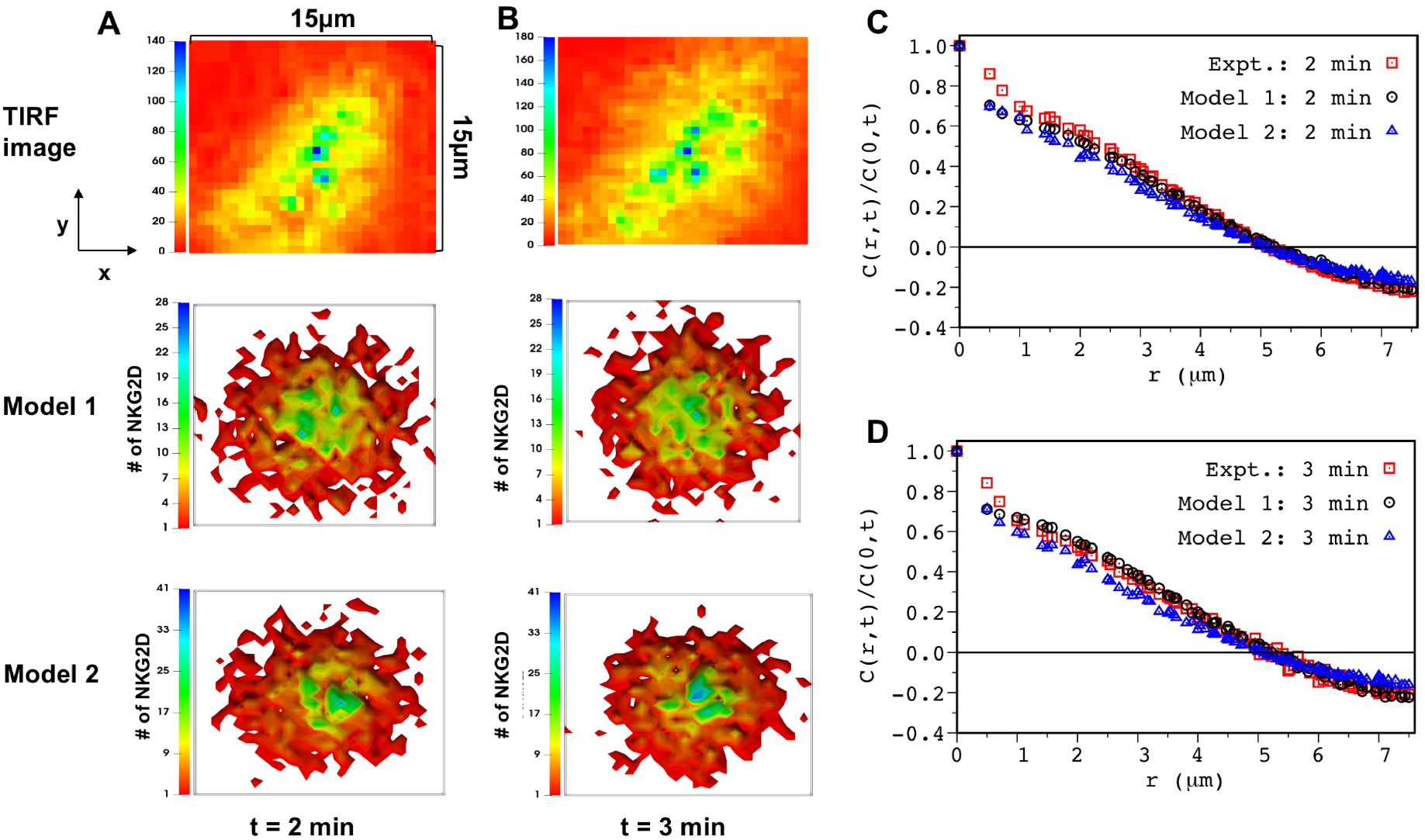
Model prediction for NKG2D spatial clustering at later times are in agreement with TIRF experiments: **(A)** (Top) Shows coarse-grained 2D image extracted for a region of interest in TIRF image (Figure S4 in Ref. ^6^) of NKG2D-Dap10-mCherry at t=2 min post stimulation by ULPB3. (Middle and Bottom) Shows spatial distribution of NKG2D in the x-y plane at t=2 min post stimulation for Model 1 (middle) and Model 2 (bottom). The model parameters are set to the best fit values obtained for the fit at t=1 min. **(B)** (top) Similar to (A), TIRF image Ref. ^6^ at 3 min post ULBP3 stimulation. (Middle and Bottom) Shows spatial distribution of NKG2D in the x-y plane at t=3 min post stimulation for Model 1 (middle) and Model 2 (bottom). The model parameters are set to the best fit values obtained for the fit at t=1 min. (C) Comparison of the two-point correlation function (C(r,t)/C(0,t) vs r) evaluated from the TIRF image (red, empty square) and simulations for Model 1 (black circle) and Model 2 (blue triangle) at t= 2 min. (D) Similar to (C) showing comparison between experiments and Model 1 and Model 2 at t=3 min.

### 2. Co-clustering of NKG2D and Vav1 is required to increase the production of pVav1 due to the formation of NKG2D microclusters

We investigated the mechanistic role of formation and centripetal movement of NKG2D microclusters in increasing pVav1 production. The average lifetime of a NKG2D-ULBP3 complex within NKG2D microclusters can increase compared to a spatially isolated counterpart because of the increase in the frequency of ULBP3 rebinding due to higher density of NKG2D molecules in microclusters. The increased ULBP3 rebinding can lead to increases in abundances of NKG2D-ULBP3 complexes, and subsequently to an increase in pVav1 production in Model 1 and Model 2. In Model 2, increase in reaction propensities due to co-clustering of NKG2D and Vav1 can additionally enhance pVav1 production. In order to evaluate the roles of ULBP3 rebinding and NKG2D and Vav1 co-clustering in increasing pVav1 production we compared kinetics of pVav1 production in Model 1 and Model 2 under two conditions: Case A: NKG2D is not allowed to form microclusters and perform centripetal movements, and, Vav1 does not co-cluster with NKG2D in Model 2. Case B: Spatial aggregations of NKG2D and Vav1 occur according to the model rules. The simulations for Case A result in spatially homogeneous distribution of NKG2D molecules in both models. Our simulations for Model 1 show that abundances of pVav1 at t=1 min for a range of ULBP3 doses have negligible differences between Case A and B (Figure 4A). The kinetics of pVav1 production for a particular ULBP3 dose also shows almost no difference between Case A and B (Figure S4A). In contrast, pVav1 abundances decrease in Case A compared to Case B in Model 2 for a range of ULBP3 doses (Figure 4B and Figure S4B). The decrease in pVav1 in Case A relative to Case B increases with the increasing ULBP3 dose (Figure 4B). These results suggest that co-clustering of NKG2D and Vav1 in Model 2 could be important for increased pVav1 production in the model.

**Figure 4.**
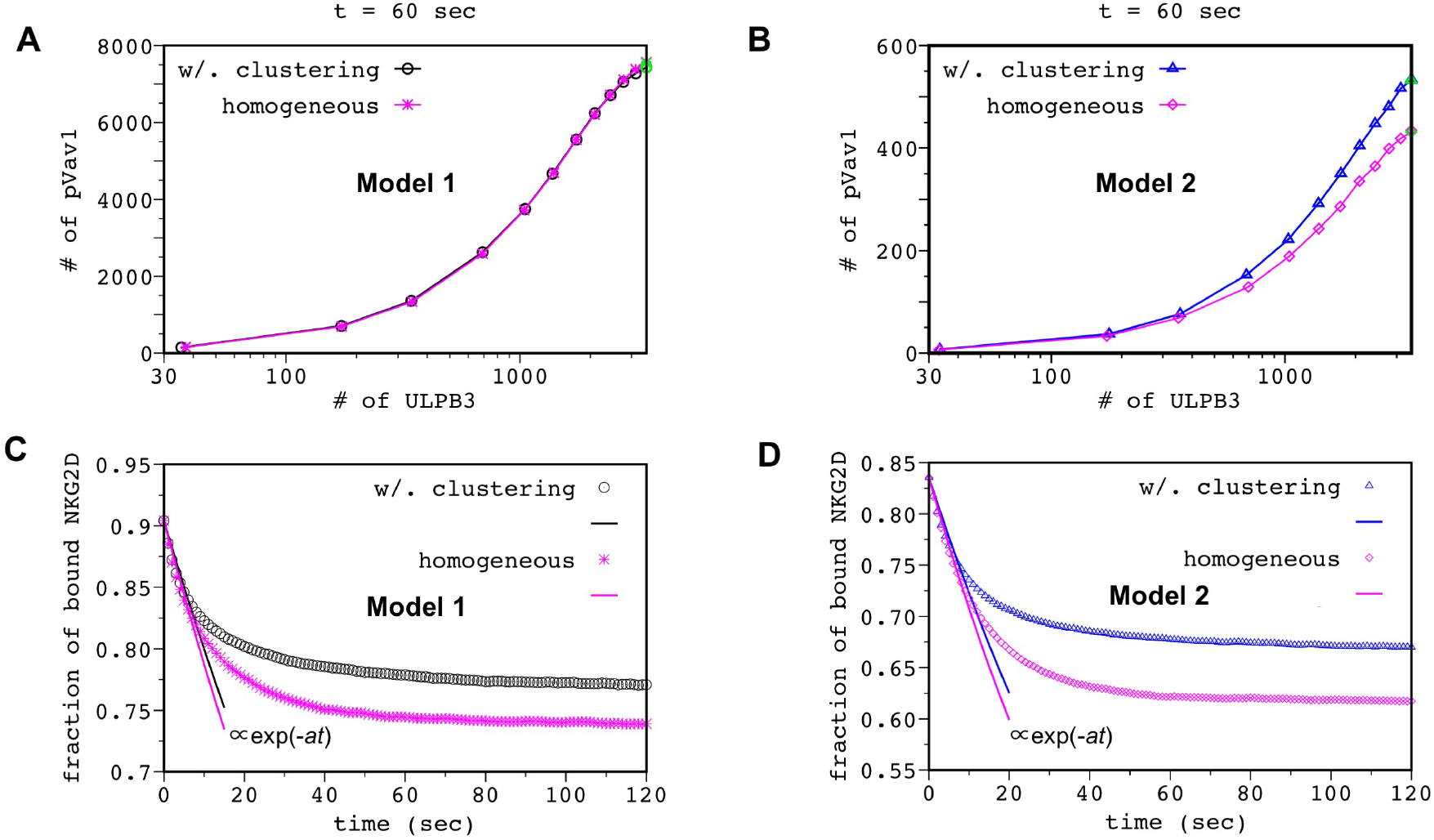
Co-clustering of Vav1 is needed to increase pVav1 upon NKG2D stimulation: Shows increase in the total number of pVav1 at t=1 min as the number of ULBP3 is increased in Model 1 **(A)** and Model 2 **(B)** for cases where NKG2D are not allowed to form microclusters (magenta asterisks, Model 1; magenta diamonds, Model 2) or form microclusters according to the model rules. The pVav1 concentrations are averaged over 50 different configurations. The points in green indicate the data obtained for parameter values that best-fit TIRF imaging data at t=1 min. **(C and D)** Decay kinetics of the fraction of the ULBP3-NKG2D complex when NKG2D molecules are distributed in space uniformly (magenta asterisks, Model 1; magenta diamonds, Model 2) or in clusters obtained from NKG2D configuration at t=1min for (C) Model 1 and (D) Model 2. NKG2D molecules are drawn from a uniform random distribution for creating the homogeneous configurations where a 0.90 (0.83) fraction of NKG2D are bound to ULBP3 at t=0 for Model 1 (Model 2). The decay kinetics are averaged over 200 initial configurations. The solid lines show fits to exponential decays at early times. The decay kinetics start deviating from the exponential decay at t ≳ 10sec.

Since ULBP3 rebinding could also play a role in increasing pVav1 production, we further quantified its contribution in increasing the average number of NKG2D-ULBP3 complexes in the Model 2. We followed an approach in ref.^33^ for this quantification, wherein decay kinetics of an initial fixed number of receptor-ligand complex is studied in the presence of immobile spatially distributed receptors. The simulations start with a fraction of receptors bound to ligands and no free ligands; free ligands created by dissociation of the receptor-ligand complex at a rate k_off_ diffuse and can rebind to the receptors. In the absence of any ligand rebinding, the number of receptor-ligand complex decays exponentially as ∝ exp(-k_off_ t). The presence of rebinding produces a non-exponential decay of the number of receptor ligand complex with higher numbers of receptor-ligand complex remaining in the system at longer times (t≫ 1/k_off_) compared to an exponential decay as exp(-*k*_off_ *t*). We evaluated decay kinetics of an initial number of NKG2D-ULBP3 complex where NKG2D-ULBP3 binding-unbinding reactions were the only reactions present in simulations. The NKG2D molecules, held fixed in space, were distributed uniformly and randomly or in a spatially clustered configuration obtained from our simulations for Model 1 or Model 2 at t= 1 min (Figure 2C). The decay kinetics shows non-exponential decay for both the cases and a small increase in the number of NKG2D-ULBP3 complex (Figures 4C-D) when NKG2D are clustered demonstrating that the effect of ULBP3 rebinding is almost equally strong when NKG2D are distributed uniformly randomly or in microclusters. Thus, increased ULBP3 rebinding in NKG2D microclusters does not play a substantial role in increasing pVav1 production.

The increase in the production of pVav1 by NKG2D microclusters in Model 2 is also consistent with experiments by Endt et al.^11^, where blocking cytoskeletal movements by an actin polymerization inhibiting drug cytochalasin D produces a substantial decrease in pVav1 in NKLs stimulated by NKG2D ligand MICA. Actin polymerization induces microcluster formation and centripetal movements of NKG2D as treatment with actin depolymerization agent latrunculin abrogated NKG2D microcluster formation and movements in NKLs^6^. Next, we used Model 2 for deciphering mechanisms of signal integration.

### 3. pVav1 dependent centripetal movements of NKG2D are abrogated by inhibitory KIR2DL2 signaling

We investigated inhibition of NKG2D signaling by inhibitory KIR2DL2 receptors in Model 2. The KIR2DL2 receptors were distributed in the simulation box following TIRF imaging data in ref. ^6^(Figure S5A). The two-point correlation function calculated from spatial distribution of KIR2DL2 at t = 0 in our simulations agrees well with that of TIRF imaging data (Figure S5C). The TIRF experiments show that KIR2DL2 microclusters are present in higher numbers at the periphery compared to the central region of the IS. In addition, the spatial organization of these microclusters does not change appreciably between pre- and ∼30s post-stimulation by HLA-C ligands (Figure S5B). The changes in KIR2DL2 microcluster distributions beyond 30s occurred presumably due to rapid retraction of NK cells from the supported lipid bilayer. Since we did not model NK cell retraction in our models, we assumed KIR2DL2 microclusters to be stationary in our simulations. The chambers where the number of KIR2DL2 exceeded a threshold value were considered to be parts of KIR2DL2 microclusters in simulations. In our models, KIR2DL2 microclusters co-localize with SFK which is supported by previous experiments^34^. NKG2D, ULPB3, HLA-C, SFK, SHP1, and Vav1 were distributed homogeneously in the simulation box at t=0. Some of the SFK molecules were co-clustered with KIR2DL2 which remained immobile for the duration of the simulations (details in Materials and Methods section). In the simulations, SHP1 recruited by the pITIMs in KIR2DL2-HLA-C complexes dephosphorylate pVav1 associated with NKG2D signaling complexes as well as free pVav1 molecules in the cytosol residing in the same spatial location or the same chamber. Thus, the production of pVav1 reduces substantially in the presence of inhibitory KIR2DL2 signaling (Figure 5B). The decrease in pVav1 abundance also decreases the centripetal velocity of the NKG2D microclusters resulting in a more spread out spatial distribution of NKG2D in simulations (Figure 5A and Movie S3). We quantified the reduction of centripetal migration of NKG2D by calculating the increase in the number of NKG2D molecules in a region enclosing the center of the IS (Figure 5C) over a time period. This decrease in the centripetal movement of NKG2D is qualitatively in agreement with experiments by Abeyweera et al. ^6^.

**Figure 5.**
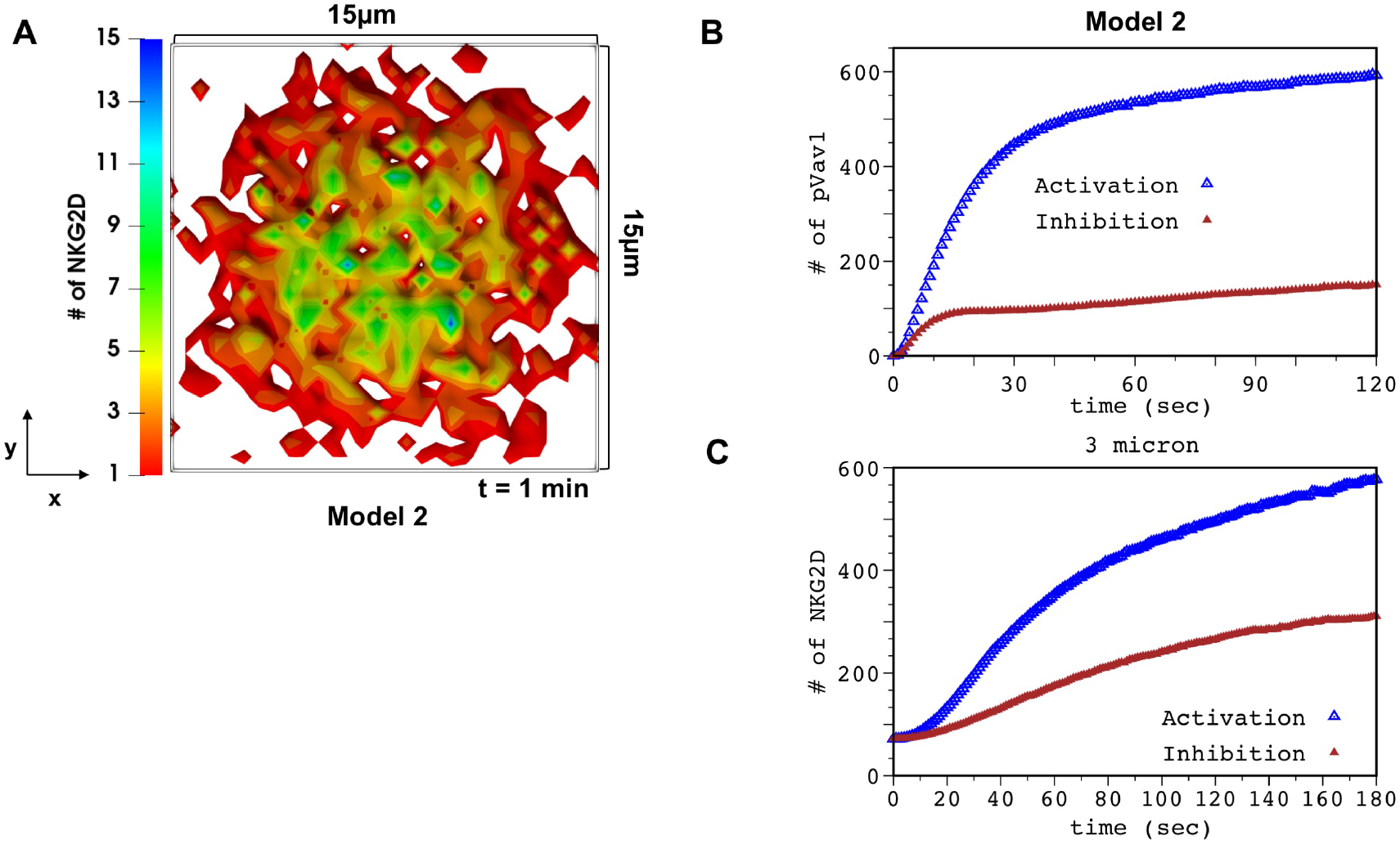
KIR2DL2 inhibition abrogates centripetal movements of NKG2D receptor clusters: **(A)** Spatial clustering of NKG2D in the presence of inhibitory KIR2DL2 signaling in Model 2 at t=1 min post ULBP3 stimulation. KIR2DL2 is stimulated by HLA-C at t=0. KIR2DL2 is distributed in the simulation box following the data extracted from TIRF imaging at 1 min 44 sec in Figure 4 of Ref. ^6^. **(B)** Kinetics of total number of pVav1 in Model 2 in the presence (filled brown triangle) and absence (empty blue triangle) of inhibitory ligands (HLA-C). **(C)** The kinetics of the number of NKG2D molecules in a region of area 3µm ×3µm around the center of the simulation box for Model 2 in the presence (filled brown triangle) and absence (empty blue triangle) of inhibitory ligands (HLA-C). The decrease in the number of NKG2D in the presence of KIR2DL2 signaling shows the decrease in the centripetal flow of the NKG2D microclusters in the simulation. The pVav1 and NKG2D kinetics shown are averaged over 200 different configurations.

### 4. Interplay between Vav1 phosophorylation and centripetal movements is required for efficient inhibition of NKG2D signaling by inhibitory KIR2DL2

We used Model 2 and Model 3 to study the role of the interplay between centripetal movements of NKG2D microclusters and pVav1 in early time integration of NKG2D and KIR2DL2 signals. In Model 3, centripetal movements of NKG2D and Vav1 are independent of pVav1 abundances, thus, the decrease in pVav1 abundances due to inhibitory signaling does not affect accumulation of NKG2D at the central region of the IS (Movie S4 and Figure 6B). We carried out simulations for two scenarios where inhibitory HLA-C ligands were distributed in the simulation box homogeneously (Case I) or in a ring pattern (Case II) (Figure 6A) devoid of HLA-C molecules at the center of the ring. Similar patterns of HLA-C on target cells have been reported by Almeida et al.^34^. Simulations were carried out with HLA-C and ULBP3 in Model 2 and Model 3. When HLA-C are distributed homogeneously (Case I), the total number of pVav1 at t=1 min post NKG2D stimulation decreases substantially in both models as the number of HLA-C increased about 4 -fold (1000 to 4000 in the simulation box) (Figure 6D). The pVav1 abundances are slightly higher in Model 3 than Model 2 owing to increased NKG2D clustering in the former model. This decrease in pVav1 with increasing HLA-C in the models is qualitatively similar to the large decrease in the percentage of lysis of human NK cell line YTS in cytotoxicity assays reported in ref. ^34^ as the abundance of HLA-C on target cells increased about 4 -fold. Next, we investigated variation of pVav1 with increasing HLA-C dose in Model 2 and Model 3 when HLA-C molecules were distributed in a ring pattern (Case II). The simulations show smaller decrease in the number of pVav1 in Model 3 compared to Model 2 as the number of HLA-C is increased (Figure 6C). This result can be explained as follows. In Model 3, NKG2D microclusters, regardless of the pVav1 concentrations, accumulate at the center of the IS devoid of HLA-C and produce pVav1 even at high HLA-C concentrations, whereas, in Model 2 the lower amount of pVav1 at high HLA-C concentrations decreases centripetal movements of NKG2D and prevents accumulation of NKG2D at the center of the IS (Figure S6). Therefore, the HLA-C ligands are able to suppress the production of pVav1 more efficiently in Model 2 compared to Model 3.

**Figure 6.**
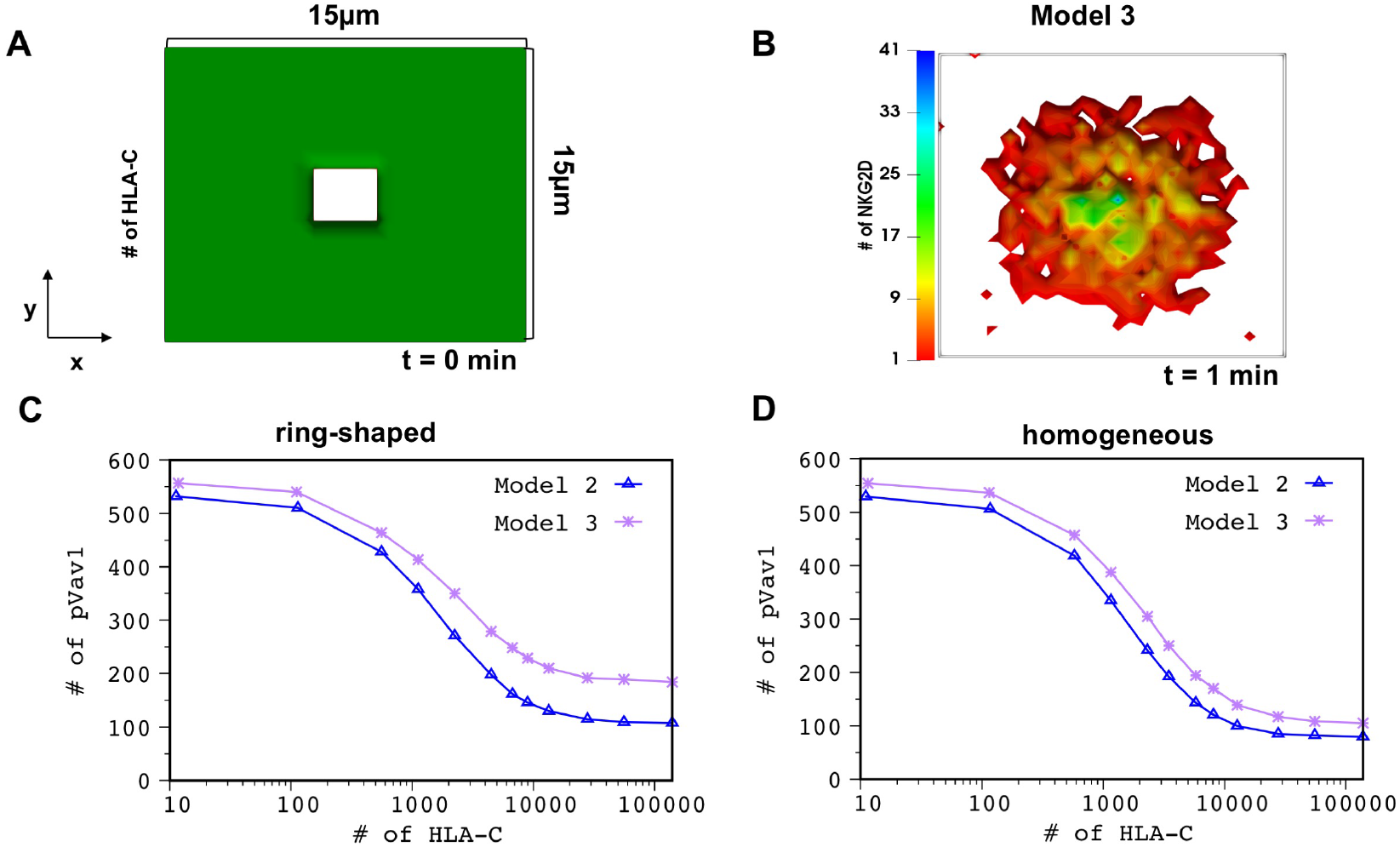
The presence of the spatial positive feedback allows for efficient inhibition: **(A)** Shows spatial arrangement of inhibitory HLA-C ligands in a ring configuration used in the simulations of Model 2 and Model 3. **(B)** Spatial distribution of NKG2D the simulation box for Model 3 at 1 min in the presence of both activating ULBP3 and inhibitory ligands shown in (A). Note the central accumulation of NKG2D because of the independence of NKG2D centripetal movements on pVav1 for Model 3. **(C and D)** Show variation of total number of pVav1 at t=1 min with increasing HLA-C concentrations for Model 2 (empty blue triangle) and Model 3 (purple asterisks). The decrease in pVav1 is higher in Model 2 compared to Model 3. The decrease in pVav1 Model 2 is more pronounced at larger HLA-C concentrations when HLA-C are organized in a ring pattern (C) vs distributed homogeneously (D). The pVav1 values shown in (C) and (D) were obtained by averaging over 50 configurations for each HLA-C dose.

## Discussion

Spatial organization of activating human NKRs such as CD16^35^, NKp46^36^, KIR2DS1^37^, and NKG2D^5-7^, and associated signaling proteins has been observed in NK cells stimulated by cognate ligands. Microcluster formation by NKRs can increase the apparent lifetime of NKR-ligand complexes because of increased frequency of ligand rebinding within a microcluster, and co-clustering of NKRs with signaling proteins can increase biochemical propensities of associated signaling reactions due to increased local concentrations of these molecules. Either of the above mechanisms can lead to an increase in the production of activated (e.g., tyrosine phosphorylated) signaling proteins. However, when the contribution due to ligand rebinding is not substantial, co-clustering of NKR and other signaling proteins in NKR microclusters plays a dominant role in increasing downstream signaling. We developed a predictive in silico framework to quantify relative roles of ligand rebinding and co-clustering of NKRs and signaling proteins for early time (∼ few mins) NKG2D signaling to demonstrate that co-clustering of Vav1 with NKG2D plays a more dominant role over ULBP3 rebinding in increasing activating signals generated by NKG2D. Since NKG2D binds with ULPB3 with a half-life (ln(2)/k_off_) of ∼ 30 seconds and the average number of NKG2D molecules in the area of a typical NKG2D microcluster when NKG2D are distributed homogeneously is about two molecules, increasing the number of NKG2D in microclusters does not result in a substantial increase in the number of NKG2D-ULBP3 complex due to ligand rebinding. Our simulations confirm the above explanation. Co-clustering of signaling proteins could play an important role for increasing downstream signaling for other activating NKRs as well. The density of CD16 on primary NK cells is about 200 times larger than that of NKG2D^38^ and the lifetime of a CD16-ligand complex is similar (e.g., k_off_ <0.01 s^-1^ for IgG^39^) or larger than that of NKGD-ULBP3, therefore, the effect of co-clustering of signaling proteins with CD16 could be more relevant for increasing downstream signaling than ligand rebinding.

Our agent based model (Model 2) is successful in generating predictions regarding inhibition of NKG2D signaling by inhibitory KIR2DL2 signaling or by drugs inhibiting actin polymerization. The model qualitatively captures the slowing down of the NKG2D microcluster movements in the presence of inhibitory KIR2DL2 signaling (Figure 5A and 5C) and the decrease in pVav1 (Figure 5B) in response to an actin polymerization inhibiting drug cytochalasin D in NKLs stimulated by NKG2D ligand MICA^11^. The model also quantitatively predicts spatial pattern of NKG2D in TIRF imaging at later time points, not included in model training, reasonably well. The model can be further expanded to investigate mechanisms of interplay between pairs of activating and inhibitory NKRs stimulated by activating ligands and different organizations of HLA-C ligands with peptides. Adaptable spatial patterns of antigens presented by DNA origami structures have been recently used to manipulate responses in B-cells^40^ and CAR T-cells^41^. The spatial model developed here can serve as a screening tool for identifying spatial patterns of antigens optimal for generating specific lymphocyte responses.

We studied the role of the interplay between creation of NKG2D microclusters, centripetal movement of the microclusters, and early time signaling using an alternate model (Model 3) where centripetal movement of NKG2D microclusters is independent of pVav1. Our simulations show that pVav1 dependent centripetal movements allow for a more efficient suppression of pVav1 in the presence of inhibitory KIR signaling, in particular, when inhibitory ligands (HLA-C) are clustered in a ring formation in the target cell membrane. HLA-C molecules have been found to organize in ring or multifocal patterns on target cells^34^, the presence of the spatial feedback in NKG2D microclusters and pVav1 creates an efficient and flexible maneuver to inhibit signaling when inhibitory ligands are present in spatially inhomogeneous patterns.

A unique aspect of our modeling framework is estimation of model parameters that best fit spatial patterns of NKG2D measured in TIRF imaging experiments. Many parameters in our model were not measured in experiments previously or are difficult to measure because they described coarse-grained processes (e.g., parameters in describing centripetal force). In addition, reported parameter values measured under specific conditions (e.g., outside the cell) might not reflect its magnitude within the cell. Therefore, we carried out an estimation of model parameters using parallel codes that produced quantitative agreement between spatial patterns of NKG2D in simulation and imaging experiments. We used two-point correlation functions to quantify spatial organization of NKG2D microclusters which is widely used in statistical physics and materials science to quantify spatial patterns. The parameter estimation step provides the following benefits: (i) it allows models to capture unique aspects of kinetics of NKG2D microclusters, e.g., centripetal movement with decreasing mobility at the center of the IS. (ii) Provide parameter ranges where the roles of ligand rebinding and co-clustering of NKG2D and Vav1 can be compared. Since both the processes depend on parameter values, these should be compared in parameter ranges that are able to reproduce experimental data. Receptor clustering in the IS has been widely investigated in computational models in T-^42-48^, B-^49^, and NK-cells^8,10^, however, most of these models described the spatial patterns qualitatively. Some of the modeling quantitatively described receptor patterns in NK-^50^ and T-^51^ cells, however, did not couple receptor clustering with signaling kinetics. A major difficulty in parameter estimation in spatial models is computationally intensive nature of such calculations, which we address here using parallel computation. However, as we discuss below there are several areas where this framework requires further improvement.

### Limitations of our work

We did not include integrin receptor LFA1 and its ligand ICAM-1 in our models for simplicity and focused our study to NKG2D and KIR2DL1 signaling and clustering. Larger size LFA1-ICAM-1 complexes (∼40nm) segregate from the smaller sized (∼10-15nm) NKG2D-MICA or KIR2DL1 molecules due to size based exclusion interactions^52^. Usually, LFA1-ICAM1 molecules reside in the periphery of the IS surrounding NKG2D^52,53^.

However, integrin receptor signaling contributes towards NK cell signaling^54-56^ and spatial clustering of LFA-ICAM1 will likely contribute towards early time NK signaling and its regulation of NKG2D spatiotemporal signaling kinetics. The models also do not include NKG2D degradation^57^ which tends to occur later in the signaling (>15 mins)^58^ and could underlie the lower amount of phosphorylated ITAM phosphorylation in the central region in the TIRF experiments in Abeyweera et al. ^6^. Furthermore, since our modeling can describe early time (0-3 min) signaling kinetics and cannot describe changes in NKG2D spatial patterns due to spreading of NK cell surface on the glass slide at times ≥ 4min (Figure S3) or later time signaling events such as secretion of lytic granules. One the computation side, several parameters showed large error bounds implying these parameters can be changed substantially but will make small or no changes to the mean cluster sizes and two point correlation functions. A potential solution to address this challenge could be introducing weight factors for combining mean values and two-point correlation functions (or second moments) following a systematic framework such as generalized method of moments^59^.

## Materials and Methods

### Spatial kinetic Monte Carlo simulation

We used the software package SPPARKS (https://spparks.sandia.gov/) to simulate reactions and particle hopping moves described below. We chose Gibson-Bruck implementation of Gillespie algorithm^60^ within SPPARKS to perform the kinetic Monte Carlo simulation.

### Biochemical reactions

Molecules in each chamber of volume (*l*_*0*_ × *l*_*0*_ × *z*) where *l*_0_ = 0.5µm and *z= l*_*0*_ (for plasma membrane bound molecules) or *z= 2l*_*0*_ (for cytosolic molecules) are taken to be well-mixed, and stochastic biochemical reactions in individual chambers are simulated using reaction propensities listed in Table 1. There are three types of protein molecules depending on their location in the simulation box: (i) Molecules residing in the NK cell plasma membrane which include NKG2D, SFK, and, KIR2DL2. (ii) Molecules residing in the NK cell cytosol which include Vav1, and SHP1. (iii) Molecules residing in the supported lipid bilayer in TIRF experiments which include cognate NKR ligands: ULBP3, and HLA-C. The binding of NKRs and their respective ligands can occur when the binding domains of these proteins are separated by short distances d_recep-lig_ ∼ 2 nm^61^. Therefore, the volume factor (*v*_recep-lig_) used for converting the unit of binding rate (k_on_) from (µM)^-1^ s^-1^ to propensity (∝ k_on_/*v*_recep-lig_) in s^-1^ in a chamber is taken as, *v*_recep-lig_ = *l*_*0*_ × *l*_*0*_ × d_recep-lig_. Similarly, the volume factor *v*_recep-SFK_ used for binding of NK cell receptors with SFKs is set to, *v*_recep-SFK_ *= l*_*0*_ × *l*_*0*_ × d_recep-SFK_, where d_recep-SFK_ ∼ 10 nm^14^. The volume factor *v*_cytosol_ for cytosolic molecules binding with NK cell plasma membrane bound complexes is taken as, *v*_cytosol_ = (*l*_*0*_×*l*_*0*_×2*l*_*0*_). In Model 2, the volume factor for binding of plasma membrane associated Vav1 molecules with other plasma membrane residing molecules is taken as, *v*Vav1-plasma = (*l0*×*l0*×*l0*).

### Microcluster formation

A pVav1 dependent processes is implemented to generate microclusters of NKG2D. A “potential” function *E*_*i*_ is associated with any chamber *i*, is given by,

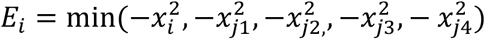

where, *x*_*i*_ is the number of pVav1 molecules in the *i*^th^ chamber and {*j1, j2, j3, j4*} denote its four nearest neighbors.

The probability of moving NKG2D molecules from the *i*^th^ chamber to its nearest neighbor *j* is given by, 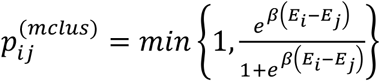; thus, when a nearest neighbor *j* has a larger potential than the *i*^th^ chamber, i.e., *E*_*j*_ > *E*_*i*_, NKG2D molecules move to the *j*^th^ chamber. We estimate *β* using PSO.

### Diffusive movements

Unbound molecules of NKRs, their cognate ligands, SFK, SHP1, and Vav1 move diffusively. The diffusive movements are simulated by hopping moves of molecules to the nearest neighboring chambers with probability p^diffu^ ∝ *D*/(*l*_0_)^2^. The propensities for these movements are given in Table 1. Periodic boundary conditions in the x-y plane are chosen for freely diffusing molecules. The propensity for diffusive movements of cytosolic molecules is about >1000 fold larger than that of the plasma membrane bound molecules and that of the majority of reactions due to the larger value of the diffusion constant (∼10 µm^2^/s)^62^ and the larger average number of cytosolic molecules (∼ 30 - 200 molecules) in a chamber. Therefore, a substantial proportion of the Monte Carlo moves are spent on diffusive moves of cytosolic molecules which make the run time of the simulation long (wall time ∼10 hrs on a single 3.0 GHz AMD EPYC processor). We approximated these fast diffusive moves by not executing explicit diffusion moves for the cytosolic molecules but homogenizing cytosolic molecules in the simulation box at time intervals proportional to the average time a cytosolic molecule will need to diffuse the length of the simulation box. We checked the validity of this approximation by comparing simulations incorporating the above approximation with those containing explicit diffusive movements for cytosolic molecules. We did not find appreciable differences between the two (Figure S7).

### Microcluster movements

Any ULBP3 bound NKG2D receptor complex is assumed to be a part of a microcluster. All molecules of a specific ULBP3-NKG2D complex, e.g., ULBP3-NKG2D-SFK, in a chamber are moved to one of the four nearest neighboring chambers in a single microcluster hopping move. The probabilities of the hops to the neighboring chambers are given by 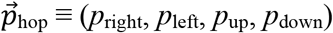 and the corresponding propensities are calculated by multiplying the probabilities by a rate 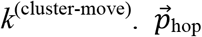 at a position (x,y) in the x-y plane is given by,

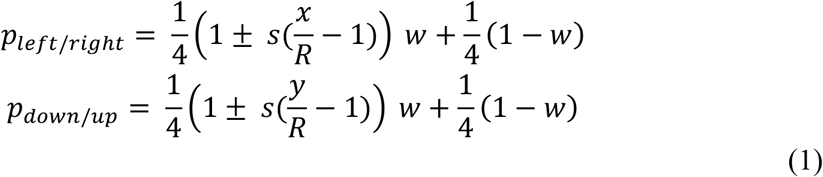

where, the four corners and the center of the simulation box in the x-y plane are at (0, 0), (0, 2R), (2R,0), (2R, 2R), and (R,R), respectively. The variable *s* depends on the total number of pVav1 (or [pVav1]_T_) in the simulation box and is given by,

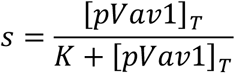

The hopping probability 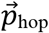 is composed of two parts: one generates centripetal movements^63,64^ and the other produces random Brownian movements. The weight factor 0≤ *w≤* 1 determines the proportion of centripetal and random components in 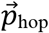, i.e., the smaller the *w*, the stronger is the bias toward random movements. The velocities for centripetal movements along x and y directions are given by,

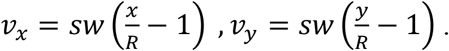

Therefore, the magnitude of the centripetal velocity *v*_r_ is given by,

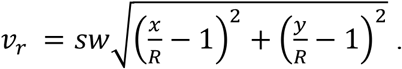

*v*_*r*_ decreases monotonically from the periphery to the center of the simulation box. *v*_*r*_ = 0 at the center (R,R) and is maximum 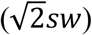 at the corners of the simulation box. In the absence of any pVav1 in the simulation box, the centripetal movements cease to exist, i.e., *v*_*r*_=0. Velocity of F-actin retrograde flow decreases monotonically from the periphery to the center in YTS NK cell lines stimulated by surface coated anti-NKG2D^12^. We assume that NKG2D microcluster movements are guided by F-actin, similar to that to TCR microclusters in antigen stimulated Jurkat T cells^65^.

The rate, *k*^(cluster-move)^ determines the magnitude of the centripetal velocity and the diffusion constant of the Brownian movements of microclusters. A movie of movements of a single microcluster is shown in the supplementary material (Movie S5).

### Excluded volume interactions

We impose an upper bound (N_thres_) to the number of molecules that can reside in a chamber to incorporate excluded volume interactions. Molecules are allowed to hop to a chamber if the number of molecules does not increase beyond N_thres_ due to the move. A different upper bound for the number of NKG2D in a chamber is implemented to prevent aggregation of NKG2D to a small number of very high density NKG2D microclusters in the central region.

### Influx of NKG2D

An influx of NKG2D molecules in the simulation box from the boundary with a rate *k*^(influx)^ is implemented by adding an outer rim of thickness 2*l*_0_ at the boundary of the simulation box. The chambers in the rim are occupied by NKG2D and ULBP3 which undergo NKG2D-ULPB3 binding-unbinding reactions. The NKG2D-ULBP3 complexes do not participate in further downstream reactions inside the outer rim. The bound NKG2D-ULPB3 complexes enter the simulation box by hopping moves that occur at *k*^(influx)^ .

### Initial configuration

Unbound molecules of NKG2D, SFK, Vav1, KIR2DL2, and SHP1 were distributed homogeneously in the simulation box where any chamber contained the same number of molecules for a particular protein. The numbers of molecules in each chamber for the above protein species are calculated using the values shown in Table 1 assuming proteins are distributed homogeneously in the plasma membrane or the cytosol of an NK cell. The number of ULBP3 in the simulation box is estimated via PSO, where, a chamber at *t*=0 is populated with 3 molecules (or left unpopulated) of ULBP3 with a probability *f* (or 1-*f*) and *f* is estimated in PSO.

### Particle Swarm Optimization

We performed asynchronous particle swarm optimization (PSO) algorithm with constraints to optimize a cost function to and estimate model parameters. The parameters were varied as powers of 10 in PSO. The parameters used in the PSO are the following: particle velocity scaling factor, *ω* = 0.5, coefficient weighting a particles best-known position, *φ*_*p*_ = 2.5, and coefficient weighting the swarms best-known position, *φ*_*p*_ = 1.5. A swarm size of 200 particles is considered. The maximum iteration limit is assigned to 100. The computation takes about 48h on 300 parallel 3.0 GHz AMD EPYC CPUs. A similar set of PSO parameters were used to estimate parameters in a spatial model describing formation of bacterial biofilms in three dimensions^66^. The construction of the cost function is described below.

### Construction of the cost function

We extracted intensity profile I({x_i_,y_i_},t) of NKG2D-DAP10 using TIRF images (details in *Extraction of image data*), where {x_i_,y_i_} denote the centers of the chambers in the x-y plane in the simulation box. Our parameter estimation scheme minimized a cost function that measures the Euclidean distance between variables quantifying statistical properties of spatial patterns of NKG2D in TIRF experiments and simulations from our agent based models. We computed mean,

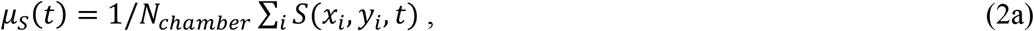

variance,

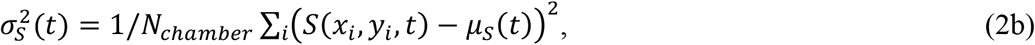

and two-point correlation function^32,66^,

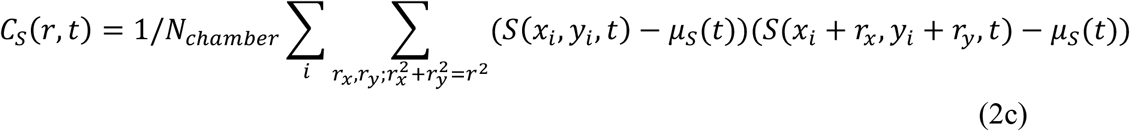

The summation over *r*_x_ and *r*_y_ indicates average of all neighbors of (*x*_i_,*y*_i_) separated by a distance *r*, i.e., 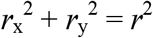. We used periodic boundary conditions for the calculation of *C*_*S*_(*r,t*) for *r* ≤ *L*/2, where *L* is the length of the simulation box. For the TIRF imaging data, *S*({*x*_i_,*y*_i_},*t*)=*I*(*x*_i_,*y*_i_,*t*)/*I*_max_(*t*), and for our model simulations, *S*({*x*_i_,*y*_*i*_}, *t*)=*n*({*x*_i_,*y*_i_}, *t*; *θ*)/*n*_max_(*t*), where *n*(*x*_i_,*y*_i_,*t*; *θ*) denotes the number of NKG2D molecules in a chamber centered at (x_i_,y_i_) in the x-y plane. *θ* represents model parameters listed in Table 1. *I*_max_(t) and *n*_max_(t) denote the maximum values of image intensity and NKG2D number in the chambers in the imaging data and in the simulation, respectively.

The quantities in Eq. (2) depend on the initial state of the system at *t*=0 in simulations or experiments and on intrinsic noise fluctuations during time evolution generating spatial configuration of molecules at time t (Figure S8). Averages of these quantities (denoted by ⟨…⟩) over ensembles of configurations at time *t* for different initial states and intrinsic noise fluctuations are traditionally used in physics^32^ to characterize spatial patterns. However, it is challenging to perform such ensemble averages in our situation because, (i) few replicates of experimental data are usually available, and (ii) computational cost of performing ensemble averaging within PSO. In order to circumvent these issues, we minimized a cost function as a function of the quantities in Eq. (2) where the best-fit values of the model parameters depend on the initial state as well as on intrinsic noise fluctuations. However, we account for effect of the intrinsic noise fluctuations in the parameter estimation as described in the error estimation section.

The cost function is given by,

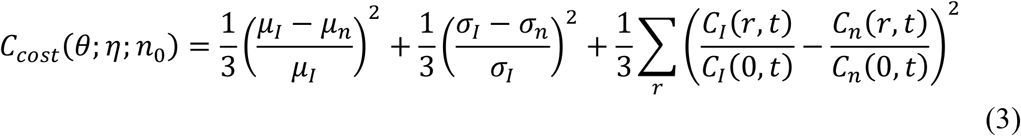

*η* represents random variables associated with intrinsic noise fluctuations and *n*_0_ denotes the initial state at t=0 in the model. The minimization of Eq. (3) in our PSO yields a *θ=θ*_*min*_ associated with a specific set of random variables *η*_pso-optim_ and a fixed initial state *n*_0_, i.e., the minimum value of C_cost_, C_cost_|_min_ = C_cost_(*θ*_min_; *η*=*η*_pso_otim_ ; *n*_0_).

### Quantification of uncertainties in PSO estimation of parameters

We generated configurations {*n*({*x*_i_,*y*_i_},*t*; *θ=θ*_min_)} for the best-fit parameter value *θ*_min_ for an initial state *n*_0_, and different intrinsic noise fluctuations (or different *η*) in our simulations. We computed C_cost_(*θ*_*min*_; *η, n*_0_) given by Eq. (3) for each *n*({*x*_i_,*y*_i_},*t*; *θ*_*min*_) and computed the variance, σ_C_, for the values of C_cost_ evaluated for the ensemble,{*n*({*x*_i_,*y*_i_},*t*; *θ*)} (Figure S9). We reasoned that the range 0≤ C_cost_≤ C_cost_|_min_ +2σ_C_, will be scanned by intrinsic noise fluctuations in the model for *θ*_*min*_; therefore, a configuration *n*({*x*_i_,*y*_i_}, *t*; *θ*) where *θ* ≠ *θ*_*min*_ that generates a cost function C_cost_ within above range cannot be separated well from {*n*({*x*_i_,*y*_i_},*t*; *θ*_min_)}. This produces an uncertainty in our estimated parameters *θ*_*min*_. We characterized the uncertainty in *θ*_*min*_ by collecting the parameters {*θ*} that produce cost functions in range 0 to C_cost_|_min_ +2σ_C_ (Figure S9). Next, we evaluated if these parameters reside in a single or multiple clusters in the manifold spanned by the parameters – existence of multiple clusters will indicate the presence of multiple local minima. We computed the number of cluster following a method in Ref. ^67^ and found a single cluster (Figure S10). Each parameter was scaled as (θ-θ_min_)/(θ_max_ – θ_min_), where θ_min_ and θ_max_ are the minimum and the maximum parameter values, respectively, to make it dimensionless and to lie between 0 and 1 before applying the algorithm in Ref. ^67^. The standard deviations computed for the points in the cluster gives an estimate of the uncertainty in *θ*_*min*_.

### Extraction ofimage data

Intensities for fluorescently labeled molecules in TIRF experiments were extracted from images published in in Ref. ^6^. The extracted intensities for fluorescently tagged NKG2D-DAP10 or KIR2DL2 molecules from regions of interest were averaged over multiple pixels to create spatial data with the minimum resolution (∼0.5 µm) in our simulation. Further details are provided in Figure S11.

### Data and code availability

Codes describing our in silico models and parameter estimation are available at the link https://github.com/jayajitdas/NK_signaling_spatial_model.

## Author Contributions

RKG and JD conceived and planned the project. RKG performed numerical simulations and calculations. RKG and JD analyzed data and wrote the manuscript.

## Acknowledgement

This work was supported by the NIH awards R01-AI 143740 and R01-AI 146581 to JD, and by the Research Institute at the Nationwide Children’s Hospital. We thank Salim I. Khakoo and C. Jayaprakash for discussions. RKG thanks John Brown, Darren Wethington, and Ali Snedden for help with simulations.

## Supplementary Material for

**Figure S1.**
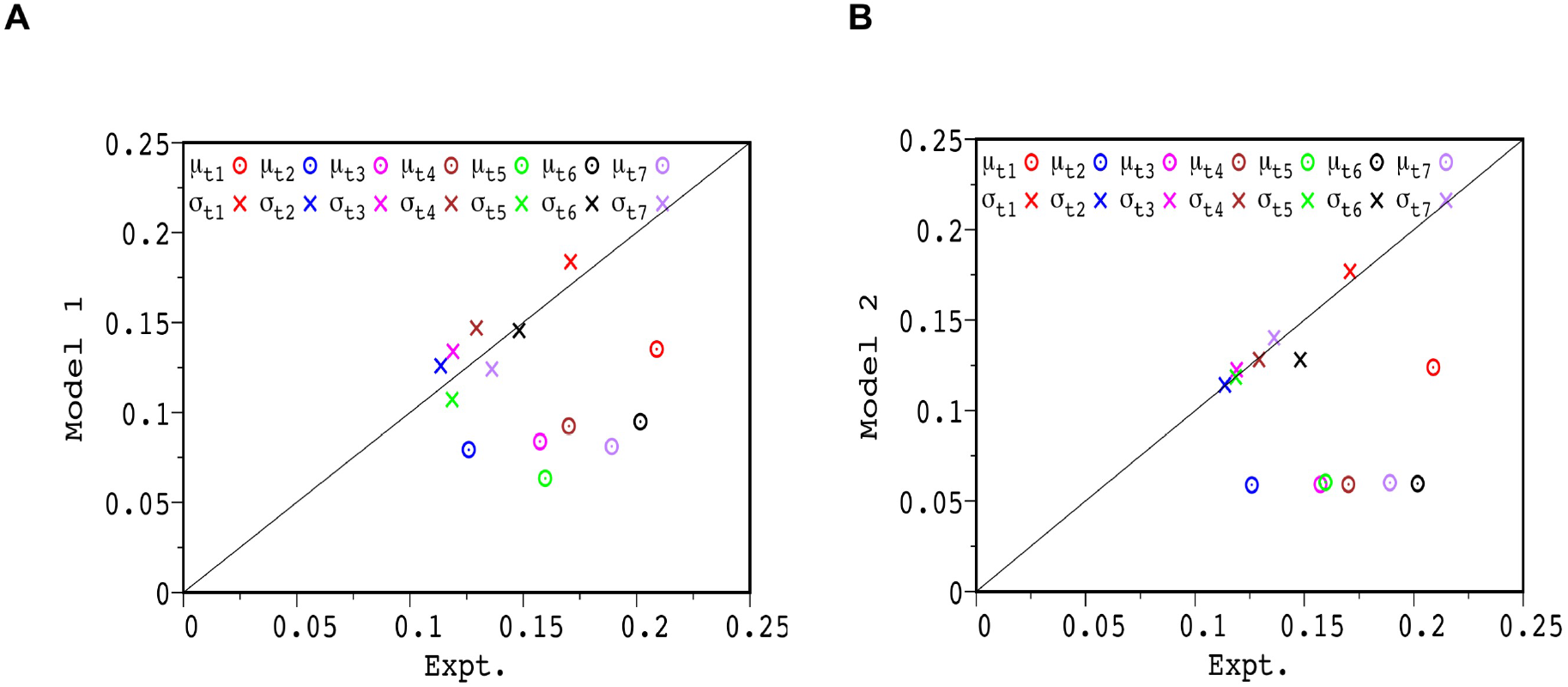
Comparison of µ_I_ and σ_I_ for TIRF images with µ_n_ and σ_n_ for configurations in model simulations. Shows µ_I_ and σ_I_ calculated from TIRF images (Figure S4 in Ref. ^1^) and model simulations at t=1, 2, 3, 4, 5, 6, and 7 mins. The calculations of µ_I_ and σ_I_ are shown in Eq. 2a and 2b in the main text. The values for TIRF images and model simulations are shown along the y and x axes, respectively. The x=y line is shown to quantify agreement/deviation between imaging data and models. The symbols indicated by µ_a_ and σ_a_ depict the values of (µ_I_, µ_n_) and (σ_I_, σ_n_) in the x-y plane at times a=t1 to t7 denoting times 1 to 7 mins, respectively. Comparisons are shown for (**A**) Model 1 and (**B**) Model 2. The models are simulated for the best-fit PSO parameters.

**Figure S2.**
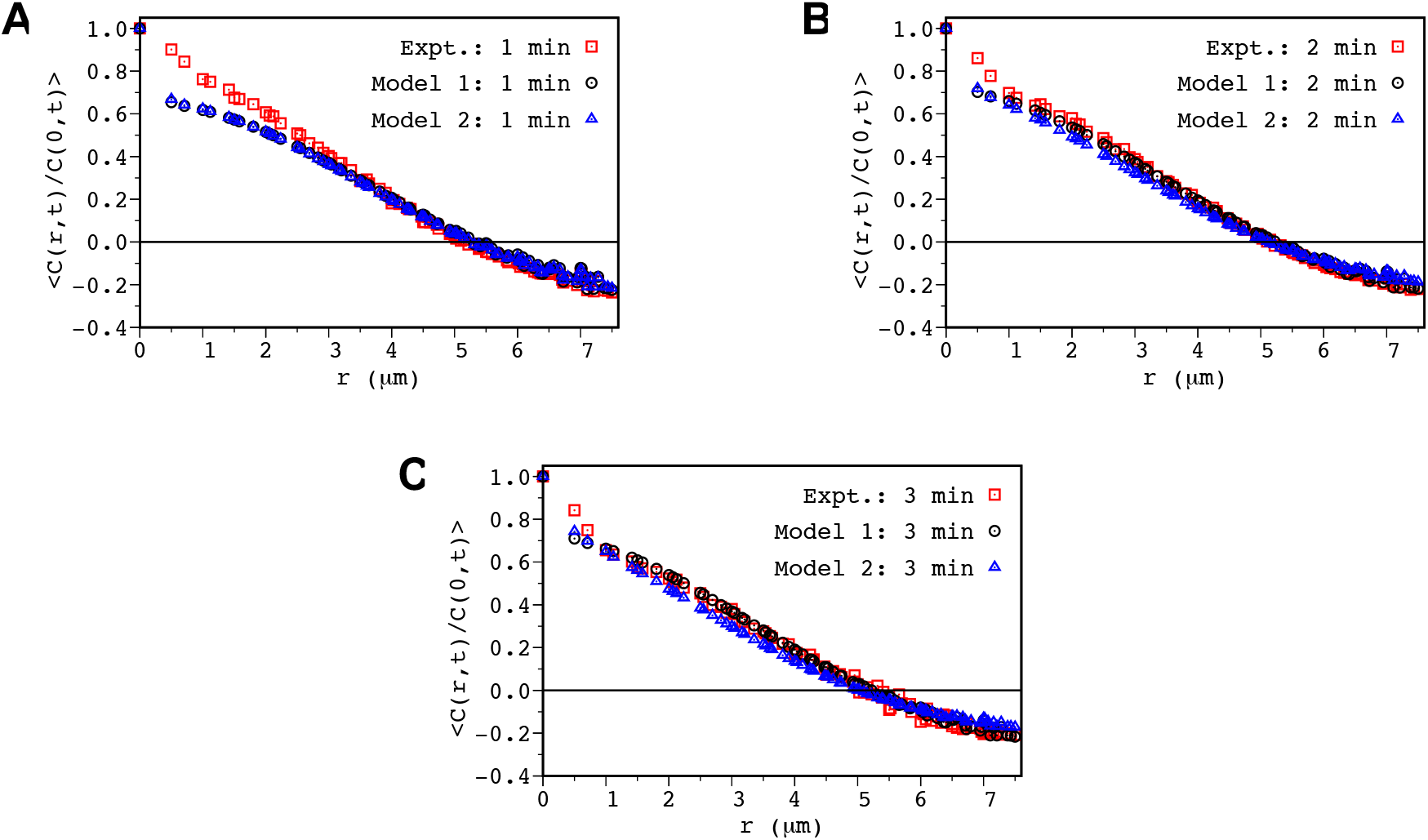
Shows comparisons between ensemble averaged two-point correlation function (<C(r,t)/C(0,t)> vs r) for Model 1 (black, empty circle) and Model 2 (blue, empty triangle) with C(r,t)/C(0,t) calculated from TIRF image (red, empty square) at (**A**) t=1min, (**B**) t=2 min, and (C) t=3 min. The parameters for the simulation are set at the best-fit values from the PSO. The two-point correlation functions for the models are averaged over an ensemble of 200 configurations.

**Figure S3.**
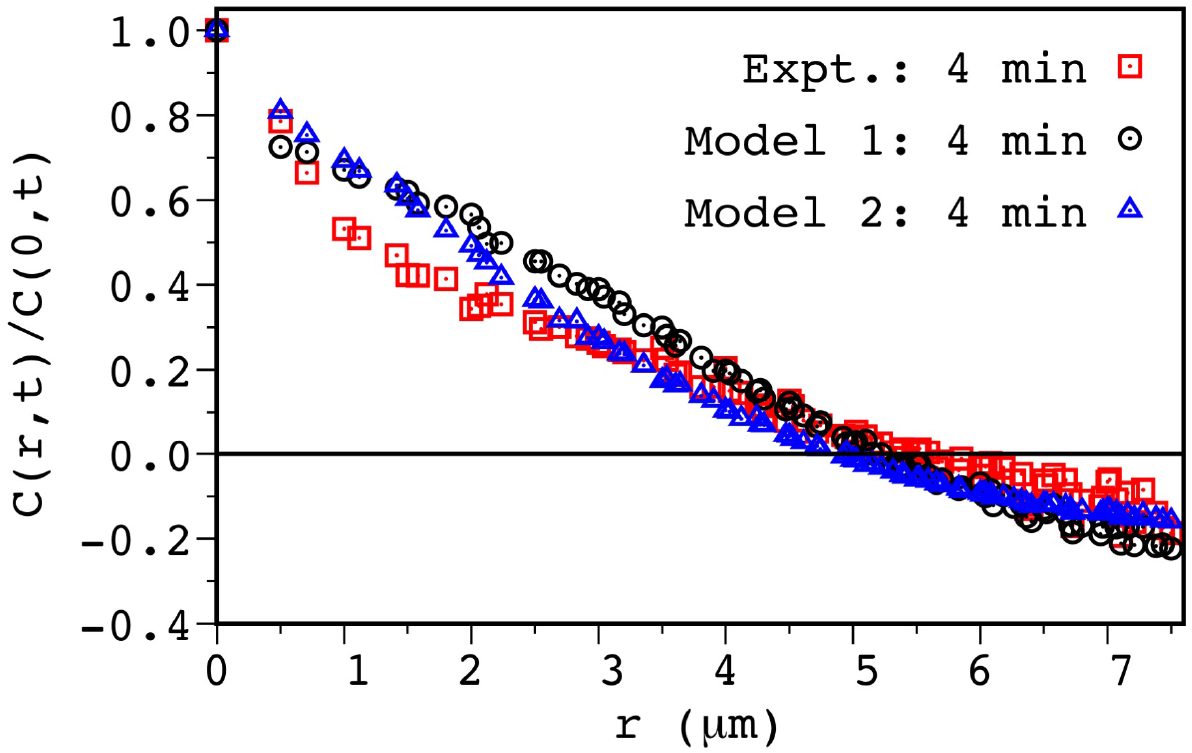
Model predictions for NKG2D spatial pattern at t=4 min show deviations from the TIRF imaging data. Shows comparison between the two-point correlation function (C(r,t)/C(0,t) vs r) at t=4 min calculated from TIRF image (red, empty square) and configurations simulated by Model 1 (black, empty circle) and Model 2 (blue, empty triangle). The parameters for the simulations are set to the best-fit values obtained from our PSO. The models show deviations from the TIRF imaging data for length scales 1-3 µm which can potentially arise due to substantial spreading of the NK cell on the lipid bilayer at 4 mins.

**Figure S4.**
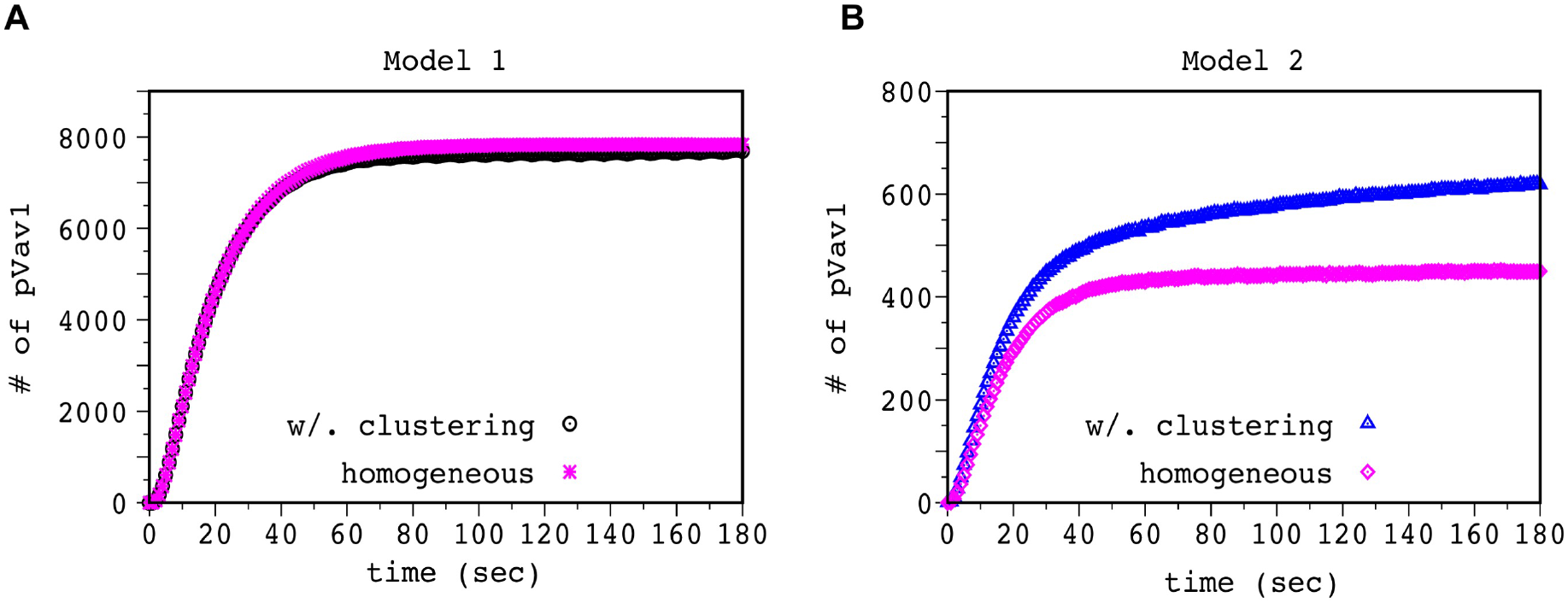
Shows increase in the total number of pVav1 with time in Model 1 (**A**) and Model 2 (**B**) when NKG2D are not allowed to form microclusters (magenta asterisks, Model 1; magenta diamonds, Model 2) or form microclusters (black open circles, Model 1; blue open triangles, Model 2) according to the model rules. The pVav1 concentrations are averaged over 200 different configurations. The models are simulated for the best-fit PSO parameters.

**Figure S5.**
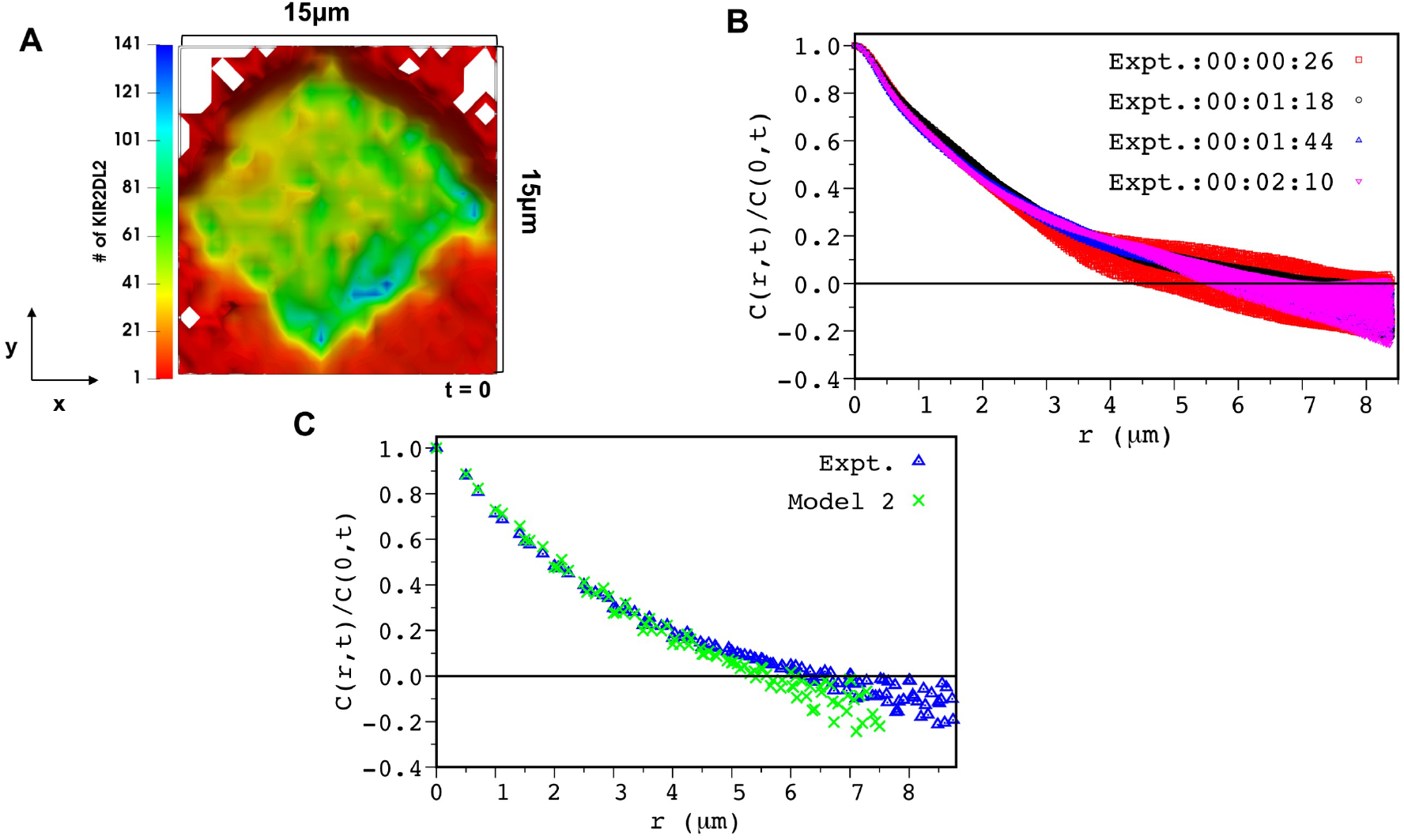
**(A)** Shows initial configuration of KIR2DL2 at t= 0 in our simulations based on the coarse-grained 2D image extracted for a region of interest in TIRF image (Figure 4 in Ref. ^1^) of KIR2DL2-GFP at t=1 min 44 sec to match the minimum length scale (∼ 0.5 µm) of spatial resolution in our model. **(B**) C(r,t)/C(0,t) calculated for intensities of KIR2DL2-GFP extracted from TIRF experiments (Figure 4 in Ref. ^1^) at times prior and ∼30s post stimulation by HLA-C ligands (UV irradiation at t = 1min 44 seconds (00:01:44), blue filled circles) for the region of interest from TIRF images. (**C**) C(r,t)/C(0,t) calculated for intensities of KIR2DL2-GFP for the region of interest from TIRF image (blue empty triangles) and after coarse-graining (corresponding to image A) to obtain the minimum scale length resolution of our simulation box (green cross).

**Figure S6.**
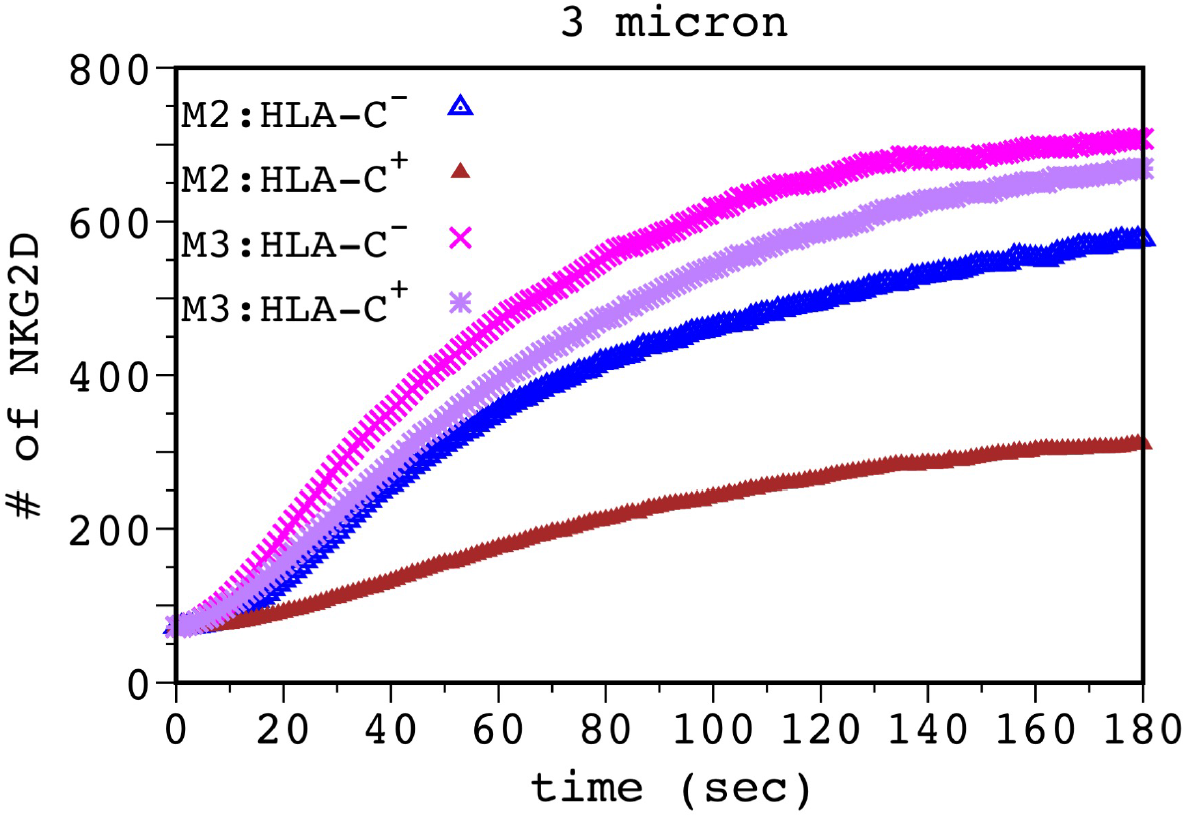
Accumulation of NKG2D in the central region of the IS in the presence of pVav1 dependent/independent centripetal motility. Shows the number of NKG2D molecules in a 3µm ×3µm area at the center of the simulation box for Model 2 (M2) and Model 3 (M3) in the presence (filled brown triangle for M2; purple asterisk for M3) and absence (empty blue triangle for M2; pink × for M3) of inhibitory ligands (HLA-C). The parameters for the simulation are set to the best-fit values. The NKG2D concentrations are averaged over 200 different configurations. KIR2DL2 inhibition abrogates centripetal movements of NKG2D receptor clusters for Model 2 but not for Model 3.

**Figure S7.**
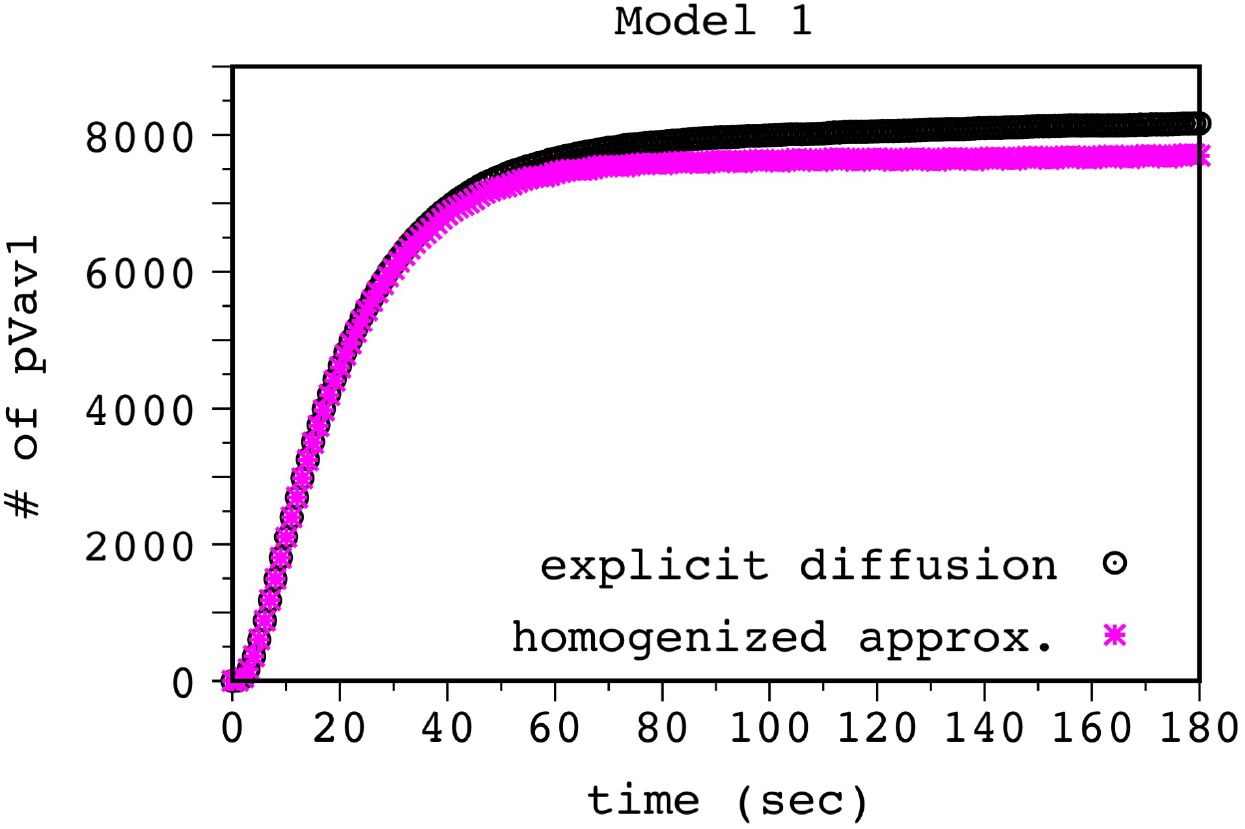
Approximating cytosolic diffusion by homogenization of molecules at discrete times has negligible effect on pVav1 kinetics for Model 1. Shows the total number of pVav1 with time in Model 1 when diffusion of cytosolic molecules is introduced explicitly in the simulations (black, empty circle), which is compared against our simulations approximating explicit diffusion with homogenization of those molecules at discrete times (magenta, asterisk). The pVav1 concentrations are averaged over 200 different configurations. The parameters for the simulation are set to the best-fit values from PSO.

**Figure S8.**
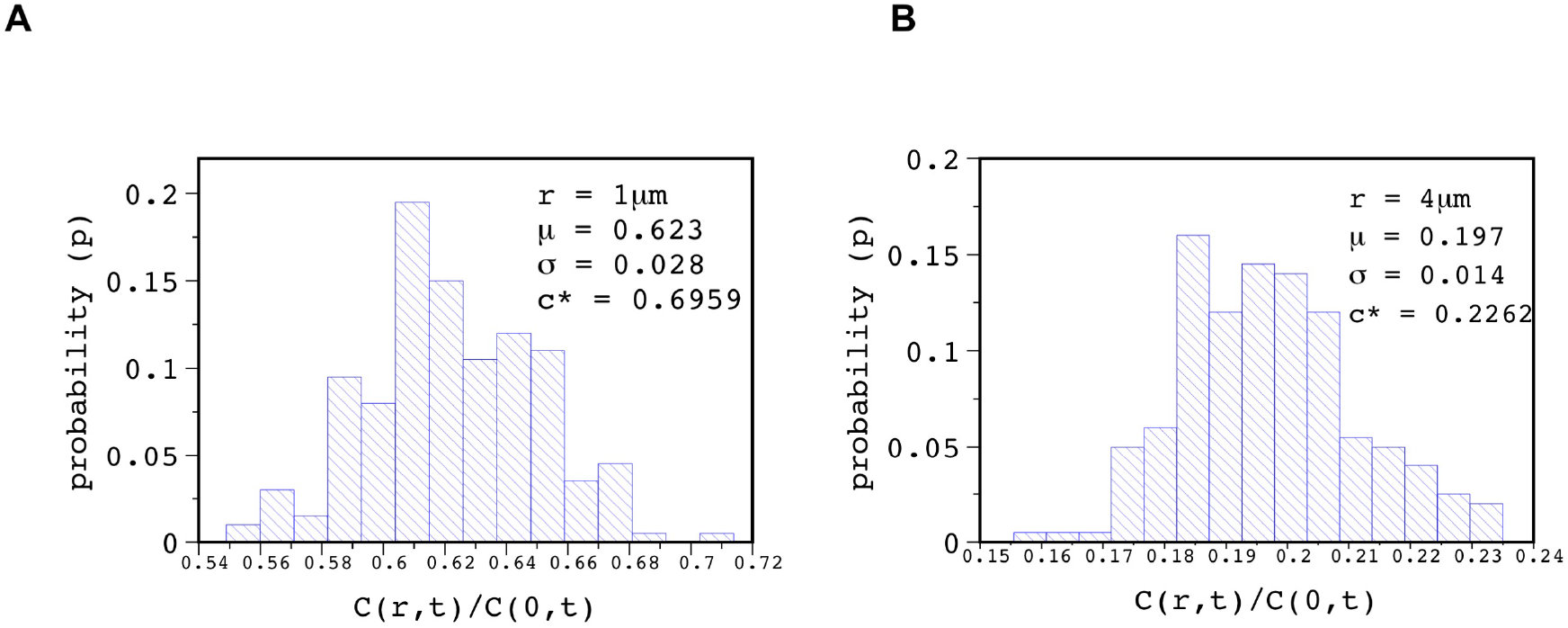
Effect of intrinsic noise fluctuations in C(r,t). Shows probability distribution function (pdf) of C(r,t)/C(0,t) at t=1min for (**A**) *r* = 1µm and (**B**) *r* = 4µm for Model 2. The parameters for the simulation are set at the best-fit value from our PSO. The pdf is calculated for an ensemble of 200 configurations. µ and σ represent the mean and standard deviation of the pdf, respectively. c* denotes the values of the C(r,t)/C(0,t) at t=1min at the r values shown in (A) and (B) for the best-fit NKG2D configuration obtained in the PSO.

**Figure S9.**
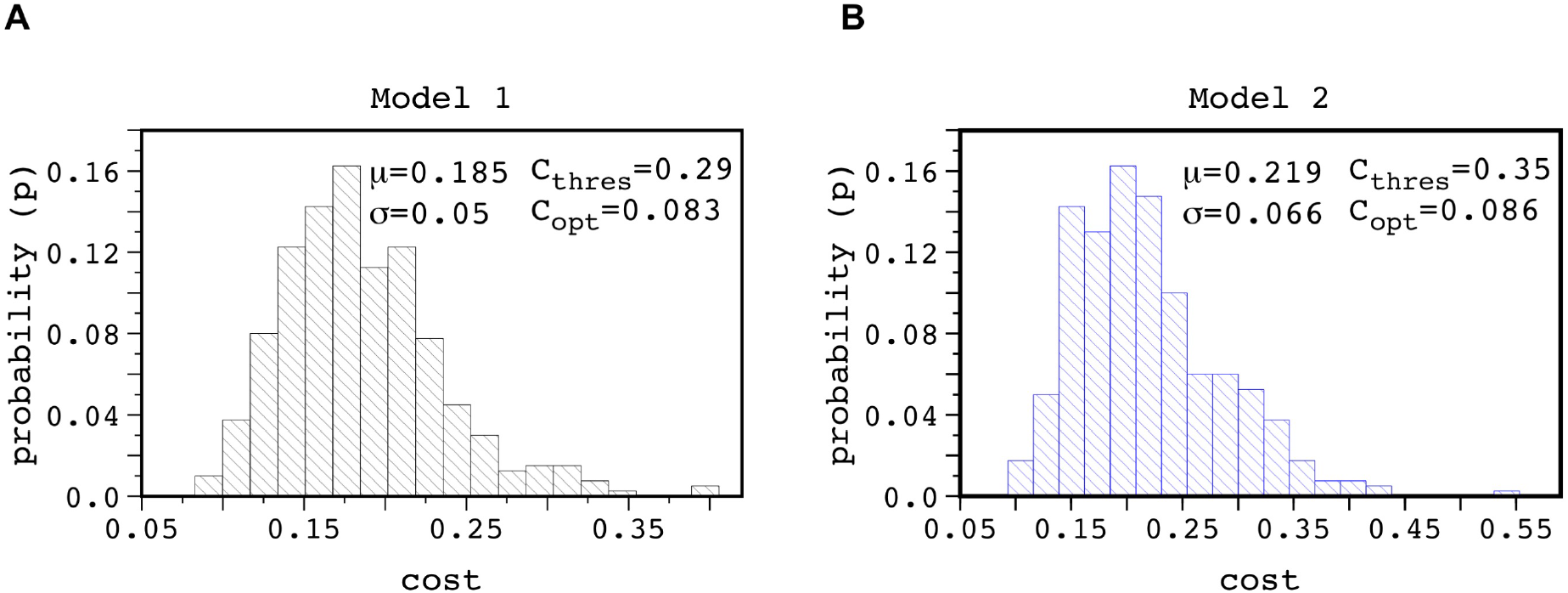
Effect of the intrinsic noise fluctuations in the cost function C_cost_(*θ*_*min*_; *η, n*_0_) for the best -fit parameter value *θ*_min_ for an initial state *n*_0_. *η* denotes the random variables associated with intrinsic noise fluctuations. The pdf is calculated for an ensemble of 400 configurations. *µ* and *σ* denotes the mean and standard deviation for C_cost_(*θ*_*min*_; *η, n*_0_) for the corresponding model. C_opt_ represents the optimum (minimum) cost function C_cost_(*θ*_min_; *η*=*η*_pso_otim_ ; *n*_0_) estimated by PSO for each model. The uncertainty in our estimated parameters *θ*_min_ is estimated to lie within the interval 0≤ C_cost_≤ C_thres,_ where C_thres_ is obtained as C_thres_ = C_opt_ +2σ.

**Figure S10.**
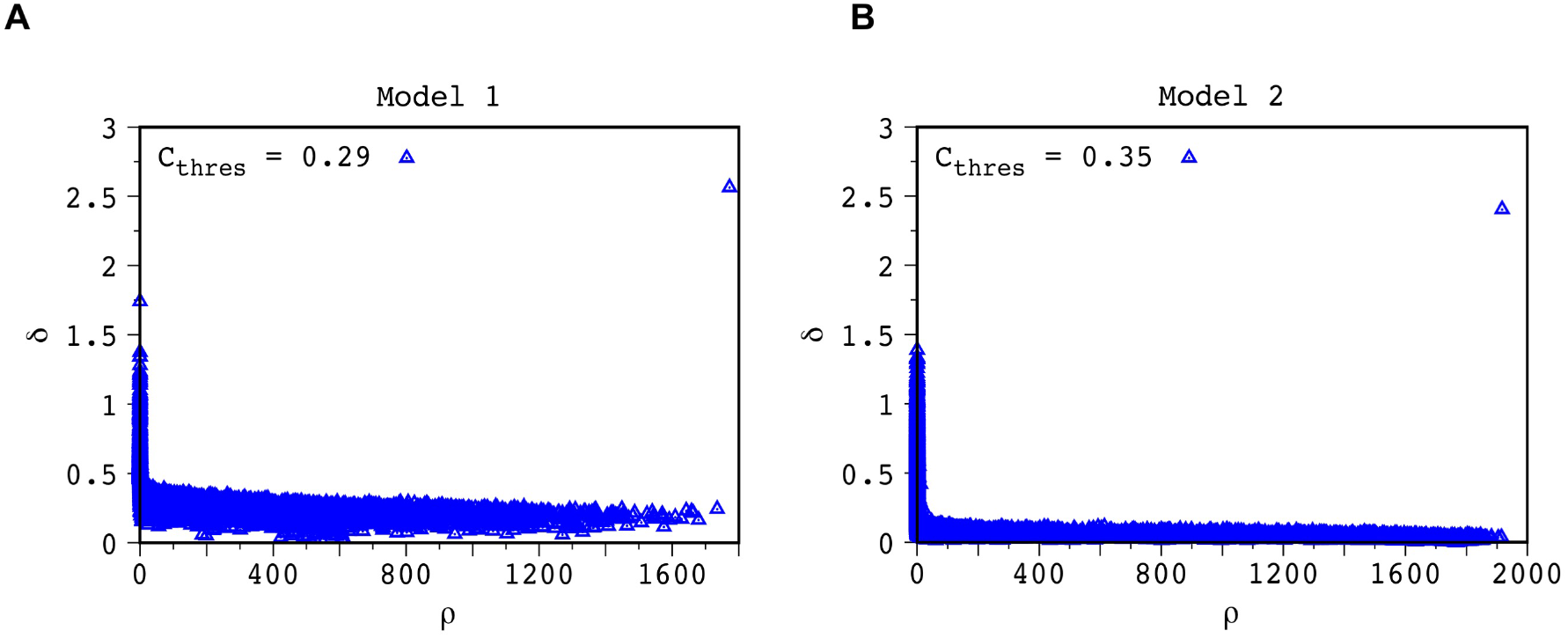
Decision Graph to compute clusters according to Density Peak Clustering algorithm (Ref. [67]). The points having relatively large local density (*ρ*) and high value of δ are considered to be the cluster centers defined in Ref. 67. For both, Model 1 (A) and Model 2 (B), we observed only one such cluster center to exist implying the presence of a single minimum cost function in the parameter range explored by the PSO. The decision graph was computed for parameters *θ* in the solution space having cost function, C_cost_ such that 0≤ C_cost_≤ C_thres,_, where C_thres_ = 0.29 and 0.35 for Model 1 and Model 2, respectively. The parameter d_c_ in the Density Peak Clustering algorithm in Ref. 67 is calculated such that the average number of neighbours is 2% of the total number of points in the data set.

**Figure S11.**
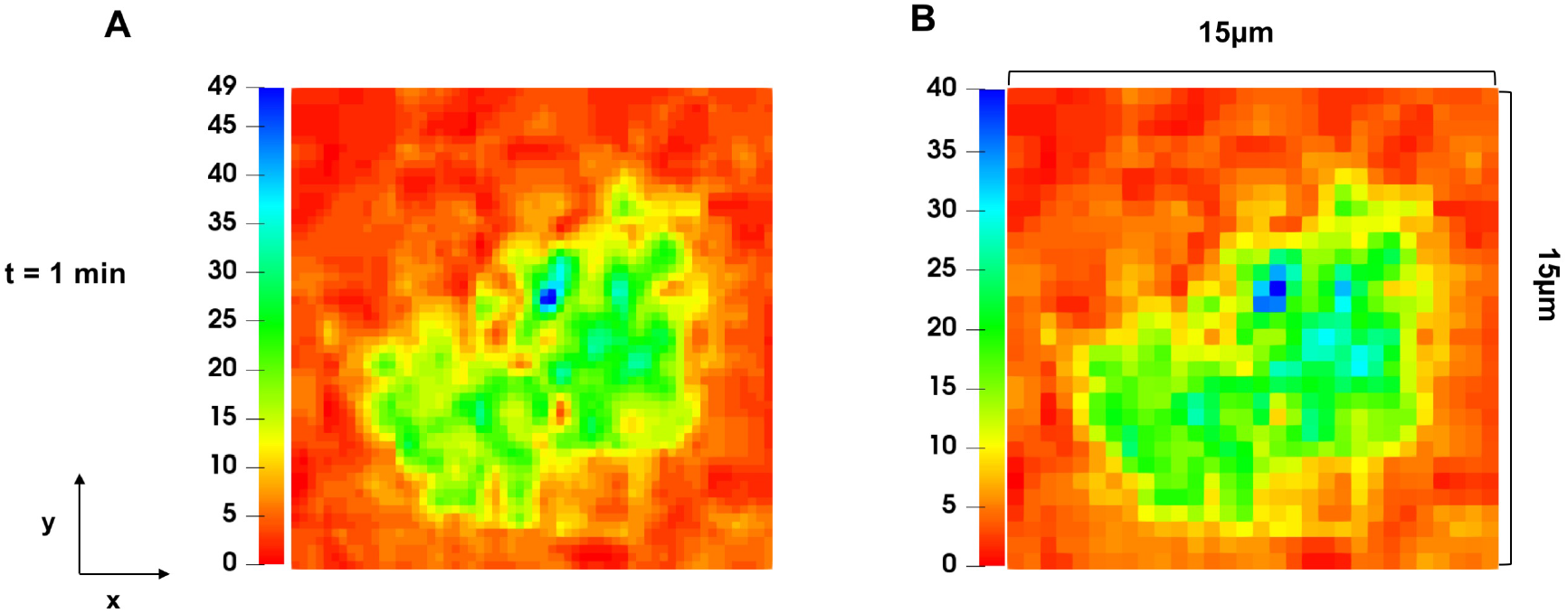
Image extraction. (A) Region of interest extracted from Figure S4 at t=1min in Ref. ^1^. (B) Coarse-grained image of region of interest in (A) to obtain the minimum scale length resolution (∼ 0.5 µm) in our simulations. The color bar shows the extracted intensities of Dap10-mCherry from the TIRF experiments in Ref. ^1^.

**Movie S1:** Shows the kinetics of spatial clustering of NKG2D in presence of its cognate ligand ULBP3 in the x-y plane in the simulation box for Model 1. The parameters for the simulation are set at the best fit value from our PSO.

**Movie S2:** Shows the spatial clustering of NKG2D in presence of its cognate ligand ULBP3 in the x-y plane in the simulation box for Model 2. The parameters for the simulation are set at the best fit value from our PSO.

**Movie S3:** Shows the decrease in clustering of NKG2D upon inhibition by KIR2DL2 in the x-y plane in the simulation box for Model 2. The parameters for the simulation are set at the best fit value from our PSO. Kinetic rates describing inhibition are provided in Table 1.

**Movie S4:** Shows clustering of NKG2D in the presence of inhibitory ligands in the absence of the pVav1 induced spatial positive feedback. The video shows clustering behavior of NKG2D in the x-y plane in the simulation box for Model 3. The parameters for the simulation are set at the best fit value from our PSO. Kinetic rates describing inhibition are provided in Table 1.

**Movie S5:** Movement of microclusters of different sizes

